# A distinct hypothalamus-habenula circuit governs risk preference

**DOI:** 10.1101/2023.01.04.522571

**Authors:** Dominik Groos, Anna Maria Reuss, Peter Rupprecht, Tevye Stachniak, Christopher Lewis, Shuting Han, Adrian Roggenbach, Oliver Sturman, Yaroslav Sych, Martin Wieckhorst, Johannes Bohacek, Theofanis Karayannis, Adriano Aguzzi, Fritjof Helmchen

## Abstract

Appropriate risk evaluation is essential for survival in complex, uncertain environments. Confronted with choosing between certain (safe) and uncertain (risky) options, animals show strong preference for either option consistently across extended time periods. How such risk preference is encoded in the brain remains elusive. A candidate region is the lateral habenula (LHb), which is prominently involved in value-guided behavior. Here, using a balanced two-alternative choice task and longitudinal two-photon calcium imaging, we identify LHb neurons with risk-preference-selective activity reflecting individual risk preference prior to action selection. By employing whole-brain anatomical tracing, multi-fiber photometry, and projection- and cell-type-specific optogenetics, we find that glutamatergic LHb projections from the medial (MH) but not lateral (LH) hypothalamus provide behavior-relevant synaptic input before action selection. Optogenetic stimulation of MH→LHb axons evoked excitatory and inhibitory postsynaptic responses, whereas LH→LHb projections were excitatory. We thus reveal functionally distinct hypothalamus-habenula circuits for risk preference in habitual economic decision-making.

## INTRODUCTION

The ability to choose the most favorable among multiple alternatives is key for an animal’s survival and reproduction in a complex, uncertain environment. When an agent confronts multiple alternatives, it calculates costs and benefits and assigns value-based preferences to all available options. Facing a decision between a certain (safe) and an uncertain (risky) option, the majority of individuals in a given population exhibit strong preference for either option which is consistent across extended periods of time^1–9^. Most individuals strongly preferred the safe over the risky option. Such prominent risk aversion has been observed in various species, ranging from birds^1^ and rodents^2, 3^ to monkeys^4–6^ and humans^6, 7^. However, once external value ratios^3, 10^ or internal state representations change ^8, 11, 12^, animals flexibly adapt their risk-sensitive choices. The similarity of risk preference-based decision-making across multiple species suggests that a shared phylogenetically conserved mechanism might be underlying. Where and how such mechanism might be implemented remains poorly understood.

In the vertebrate central nervous system, neuromodulatory regions in the mid- and hindbrain and their projection targets are known to be crucial for value-guided behaviors ^3, 13–20^. A main regulator of these regions, via strong excitatory projections, is the evolutionary conserved lateral habenula (LHb)^21–26^, which in turn receives glutamatergic and GABAergic long-range inputs from various regions in the forebrain, thereby functionally connecting the forebrain with mid- and hindbrain regions^27–31^. Whereas the LHb is mainly known for encoding negative motivational value^25^ and aversion^32, 33^, as well as for its pathological transformation in major depressive disorder^34–37^, it also has been implicated in the evaluation of economic risk. Specifically, pharmacological inactivation of the LHb in rats annihilated the ability to make economically favorable decisions when presented with a choice between different probabilistic reward outcomes^10^. However, what information is encoded in the LHb during risky decision-making, how this information is implemented, and what are the relevant synaptic inputs contributing to it remained unclear.

Here, using a balanced two-alternative choice task, we measure risk preference under risk of loss in head-fixed mice. This paradigm allows to test voluntary value-based decision-making over extended periods of time in a highly standardized manner. On the basis of this paradigm, we have used single and multi-fiber photometry, in vivo electrophysiology, longitudinal two-photon calcium imaging through chronically implanted gradient index (GRIN) lenses, and projection- and cell-type-specific optogenetics to study activity patterns in the LHb and LHb input regions, as well as behavioral adaptations during risk preference-based decision-making.

## RESULTS

### Mice display strong preferential traits during risky decision-making

To address this question, we trained head-fixed mice in a balanced two-alternative choice task. Performing this task, mice could choose between two waterspouts delivering various amounts of reward upon licking. The ‘safe’ spout delivered a reward of 5 µl sucrose water in 100% of cases whereas the ‘risky’ spout delivered either 17 µl in 25% or 1 µl in 75% of all cases (Figure 1a). Hence, the mean expected value was equal for both options. In every session, mice performed 50 forced-choice trials where in each trial only one spout (indicated by a light-emitting diode, LED) delivered a reward upon licking (with size according to safe or risky side), followed by 200-300 free-choice trials where both spouts carried a potential reward. Facing these two alternative options, most animals displayed a strong preference for either the safe or the risky option whereas only few animals frequently switched between spouts (Fig. 1b, c; Methods). In the following, we refer to these different types of individuals as ‘risk-averse’ (n = 32), ‘risk-prone’ (n = 15), and ‘risk-neutral’ mice (n = 5). Mice maintained their risk preference level (defined as the likelihood of choosing their preferred option; Methods) throughout the balanced reward sessions, even if spouts associated with safe and risky option were inverted (Extended Data Fig. 1a-c; Methods). However, they did still evaluate the available options, as they adapted their choices whenever the expected value of one option became advantageous over the other (Extended Data Fig. 1d). Moreover, risk averse and risk prone mice differed in their applied strategies coping with previous trial outcomes. Since mice mainly incorporated the previous 1-2 trial outcomes in their present decision (Fig. 1d), we calculated transition matrices (Fig. 1e) for the present choice (y-axis) depending on the previous (trial-1; x-axis) trial outcome for individual mice and pooled them according to their risk-preference. While transition matrices for risk neutral and risk prone mice were similar, they differed substantially form risk averse mice (Fig. 1e-g). For instance, after experiencing a loss, compared to risk prone, risk averse mice were more likely to switch back to the safe option or show an indecisive choice and less likely to take the risky option again. Consequently, correlation of behavioral transition matrices clearly segregated risk averse and risk prone individuals into risk preference-based clusters (Fig. 1h). Moreover, we did not detected differences between risk averse and risk prone mice based on their level of motivation, decisiveness, anxiety, or sex (Extended Data Fig. 2) which are potential confounding factors for studying risky decision-making ^38–40^.

**Fig. 1.**
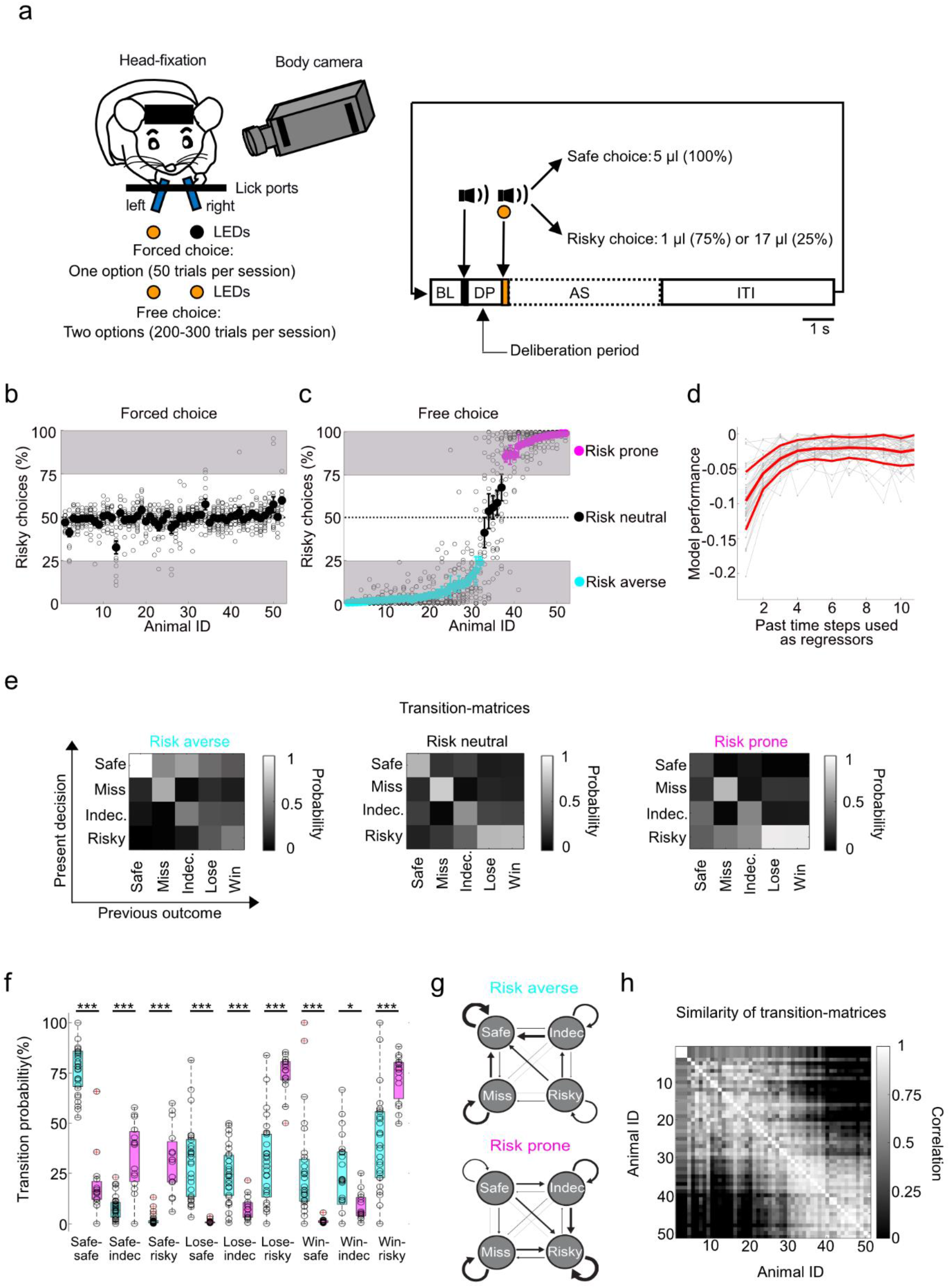
Mice display strong preferential traits for safe or risky options that are stable across time. **a**, Task structure of two-alternative choice task. After a 1-s baseline period (BL) an auditory tone indicates the start of the deliberation period (DP; arrow and black vertical bar), which terminates with the reward-indicating cues (auditory tone paired with illumination of one or two LEDs; arrow and orange vertical bar), and is followed by the action selection (AS) and the inter-trial interval (ITI) period. Fraction of risky choices during forced (**b**) and free (**c**) choice trials (n = 52 mice). Each light grey circle represents the fraction of risky choices in an individual session (n = 14 sessions for each mouse). Dark circles are mean ± s.e.m. Based on their choice preference mice were classified as risk-averse (cyan), risk-prone (magenta), or risk-neutral (black). **d**, Cross-validated multinomial linear regression was used to predict choices using past outcomes as regressors. The performance of the model was computed from the cross-entropy loss function of predictions. Grey lines indicate performances for single mice (n=52), red thick line indicates average across mice, red thin line the standard deviation across animals. **e**, Averaged transition matrices for risk-averse (left), risk neutral (middle), and risk-prone (right) mice. X-axis indicates previous trial outcome (Safe, Miss, Indecisive (Indec.), Lose, or Win) and Y-axis the choice in the current trial. Grey values represent transition probabilities. **f**, Quantification of outcome dependent trial-transition probabilities for risk averse (cyan, n=32) and risk prone (magenta, n=15; p-values from left to right: 1.12e-7; 4.98e-6; 0.8e-7; 0.8e-7; 0.74e-4; 0.95e-6; 0.41e-5; 0.027; 0.49e-4). **g**, Illustration of trial-type dependent state transition. Arrow thickness is weighted for transition probability. **h**, Correlation of individual transition matrices across mice.

### LHb population activity reflects individual risk preference

Based on the finding that pharmacological LHb inhibition prevents the formation of choice preferences in animals freely choosing between two probabilistic options^10^ (even if one option is unambiguously advantageous over the alternative), we aimed to record LHb activity during the two-alternative choice task. To monitor neuronal activity in the LHb, we expressed Cre-dependent GCaMP6f in the LHb of Vglut2-Cre mice (n = 9) and applied fiber photometry (Fig. 2a-e). Surprisingly, we found that the glutamatergic LHb population calcium signal was consistently larger for the individuals’ preferred option (be it safe or risky) prior to action selection, i.e., in the deliberation period (DP) ^41, 42^ between start and reward cue (Fig. 2f-h). This effect was abolished when we sorted fluorescence traces based on the animal’s choice in the previous trial (Fig. 2 i-k), indicating that LHb signaling in the DP reflects information related to upcoming choices rather than encoding past outcomes^3^. LHb population activity in the reward period (1-s window after valve opening) or the post-reward period (1 to 5 s after valve opening) were not able to explain risk-taking behavior of individuals to a similar extent (Extended Data Fig. 3). Furthermore, the difference in LHb activity during the DP for preferred vs. non-preferred choices could not be explained by elevated motor actions (Extended Data Fig. 4). For validation, we also extracellularly recorded action potentials in the LHb and found that multi-as well as single unit activity in the DP was higher for preferred choices, confirming that our photometry results represent increased neuronal spiking activity in the LHb (Fig. 2l-u).

**Fig. 2.**
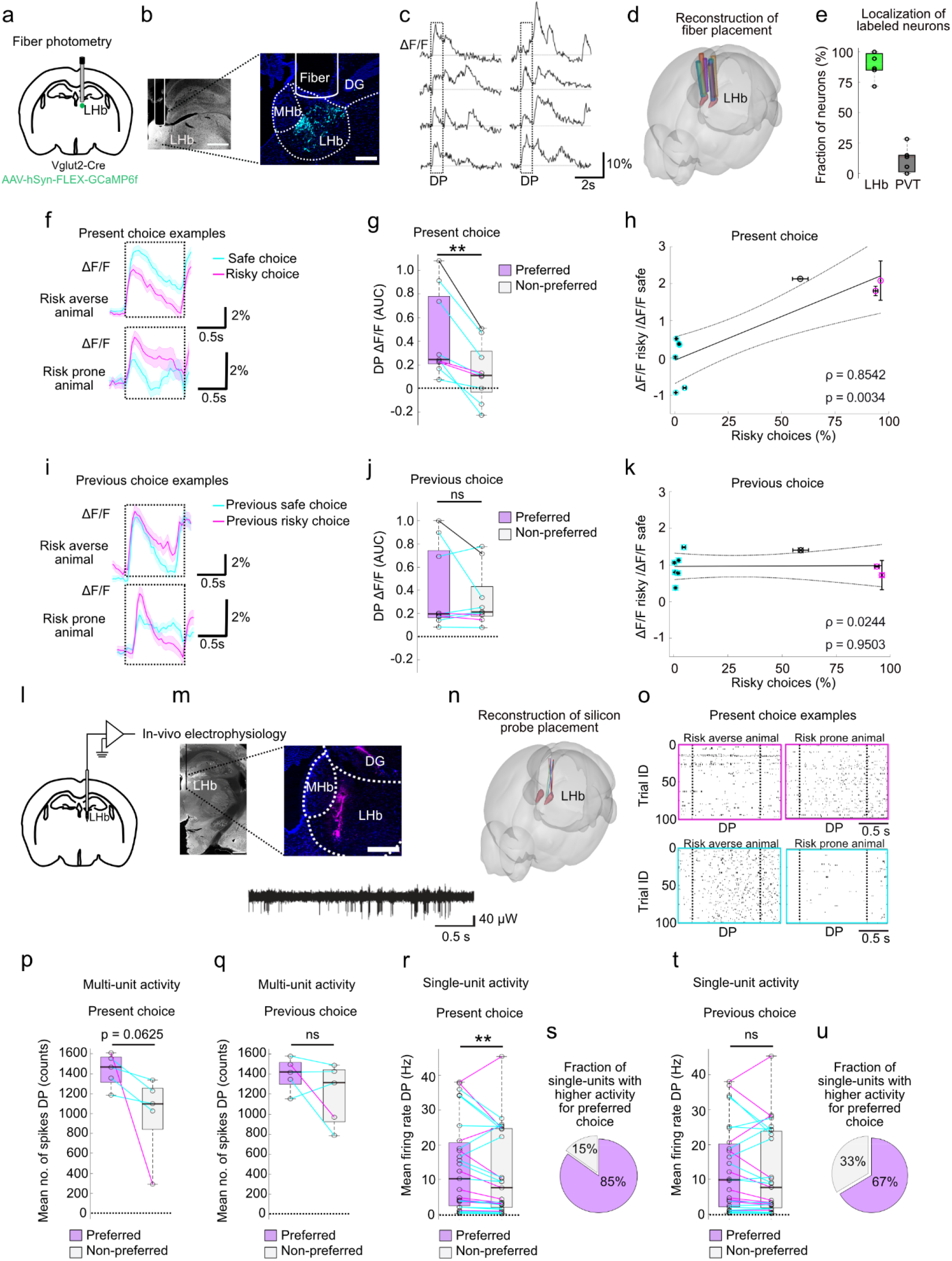
LHb population activity resembles individual risk preference prior to present action selection. **a**, Illustration of fiber-photometric recordings in the LHb. **b**, Left: Widefield fluorescence image of fiber placement in an example animal (scale bar 500 μm). Right: Confocal image of GCaMP6f-expressing LHb neurons with the fiber position indicated. Medial habenula (MHb), dentate gyrus (DG). Scale bar 200 μm. **c**, Examples of single-trial fiber-photometric calcium traces. Dashed boxes indicate DP. **d**, Reconstruction of fiber placement across all 9 mice (indicated by different colors). **e**, Localization of GCaMP6f-labeled somata (LHb and off-target paraventricular thalamic nucleus (PVT)) within the range of fiber fluorescence detection (max. 400 μm).**f**, LHb population activity during the DP (dashed box) prior to safe (cyan) and risky (magenta) choices, exemplified for a risk-averse (top) and a risk-prone mouse (bottom; 50 randomly selected trials for each condition, respectively). Data are plotted as mean ± s.e.m. **g**, Quantification of LHb population activity integrals during the DP (AUC, area under the curve) for preferred (violet) and non-preferred (grey) option (n = 9 mice). Line colors indicate individual risk preference (risk averse (cyan), risk neutral (black), risk prone (magenta)). **h**, Ratio of LHb ΔF/F integrals during the DP for present risky and safe choice population activity correlated with individual risk preference across sessions. **i-k**, Same as (f-h) but trials were sorted for choice in the preceding trial. **l**, Illustration of silicon probe recordings. **m**, Widefield fluorescence image of half hemisphere (left) and confocal image (right) illustrating probe location with example trace. Probe location is indicated by lipophilic dye (magenta). **n**, Reconstruction of silicon probe placement across all 5 mice. **o**, Single trial raster plots of detected action potentials in 100 randomly selected trials from two example animals (Right: risk averse, Left: risk prone) . Box color indicate respective trial choices (safe choice (cyan); risky choice (magenta)). Dashed lines indicate onset of start and end of reward cue and enclose the DP. Quantification of LHb multi-unit population activity during the DP for preferred (violet) and non-preferred (grey) option for present (**p**) and previous (**q**) choice trials (n = 5 mice). Single-unit activity and fraction of units with higher activity for individual preferred (violet) or non-preferred (grey) choice present (**r**, **s**) and previous choice (**t**, **u**) trials (n=27 identified single-units across 4 mice). **p<0.01, Wilcoxon signed rank test (paired). Exact p-values for all figures are provided in Extended Data Table 1.

### A subpopulation of LHb cells encodes individual risk preference

To understand how risk preference selectivity is encoded on the level of individual LHb neurons, we expressed Cre-dependent GCaMP6f in the LHb of Vglut2-Cre mice (n = 12) and imaged LHb neurons longitudinally through an implanted gradient index (GRIN) lens with two-photon microscopy (Fig. 3a-e). Across mice, most LHb neurons increased their activity during the DP compared to baseline activity, while some did either not respond or decrease their activity (Fig. 3f, Extended Data Fig. 5a). In line with our fiber photometric and silicon probe recordings (Fig. 2f-k) LHb single cell DP activity did not develop over the course of sessions (Extended Data Fig. 6) indicating that this activity does not merely reflect stimulus association learning^43^. However, independent of trial choice (risky or safe), DP activity moderately attenuated over the course of a session (Extended Data Fig. 6). This might go along with a general increase in satiation and corresponding decrease in reward valuation. Moreover confirmatory to our fiber photometry data, we found that two-photon bulk activity (acquired by drawing a single region of interest across all recorded neurons in the field of view) during the DP was higher for the preferred compared to the non-preferred option and explained behavioral performance better for present than for previous choice (Extended Data Fig. 5 b-e). This result was further confirmed on the level of individual LHb neurons (Fig. 3g). During the DP, 74.5% of recorded LHb neurons (n=409) altered their mean activity >10-fold compared to the pre-cue baseline period. Of these responsive cells, 85.1% (n=348) showed diverging activity (significant difference between preferred and non-preferred choice compared to shuffled data and receiver-operating characteristic (ROC) area under the curve (AUC) value >0.65) during the DP reflecting the animal’s subsequent choice of its preferred or non-preferred option. We refer to these cells as ‘risk-preference-selective cells’ (RPSCs). Whereas most RPSCs showed higher activity for the preferred option (47.9% of responsive cells, n=196; ‘positive’ RPSCs, blue; Fig. 3g, h), a minority of responsive cells (37.2%, n=152) displayed higher mean calcium signals for the non-preferred option (‘negative’ RPSCs, red). The ‘degree of selectivity’ (indicated by the absolute area between the curves) was further similar for positive and negative RPSCs (Fig. 3i). When we correlated the selectivity of RPSCs with their responsiveness (mean of z-scored activity during DP), we found a strong negative correlation of non-preferred choice responsiveness with selectivity for positive RPSCs (Fig. 3j, k), indicating that the degree of positive but not negative RPSC selectivity derives from the degree of DP tuning for the non-preferred choice option. Interestingly, negative RPSCs displayed a higher degree of selectivity for different reward volumes upon delivery (Extended Data Fig. 7), pointing towards different functions for positive and negative RPSCs in reward processing. RPSCs DP activity prior to choosing the risky or safe option was further closely linked to individual risk preference. Based on their selectivity, positive RPSCs displayed a potent positive correlation whereas negative RPSCs were strongly anti-correlated to behavior choices (Fig. 3l). In line with fiber photometric recordings this correlation disappeared once we sorted fluorescence traces based on the animal’s previous trial choice (Fig 3m), leading us to conclude that RPSC activity encodes information for upcoming choices rather than past outcomes.

**Fig. 3.**
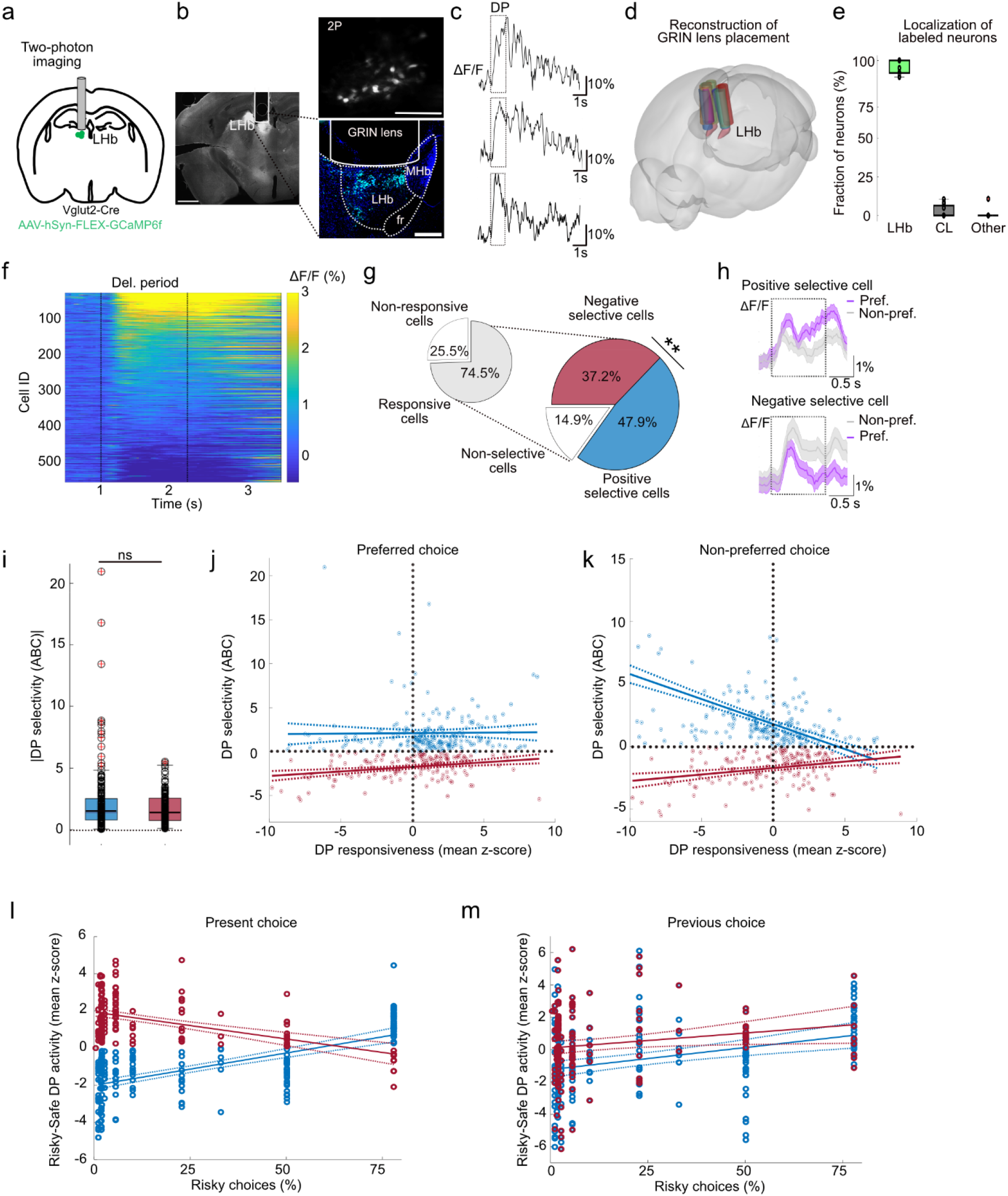
Selective LHb cells encode individual risk preference prior to action selection. **a**, Illustration of two-photon imaging through a GRIN lens (n = 12 mice). **b**, Left: Widefield fluorescence image of half hemisphere, Right: Two-photon image through GRIN lens (top) and confocal image (bottom) of GCaMP6f-expressing LHb neurons with the lens position indicated. Scale bar 1 mm (Widefield), 200 μm (2P and confocal). **c**, Single-trial traces (smoothed) from three example neurons from the mouse shown in b. Box indicates DP. **d**, Reconstruction of GRIN lens placement across all 12 mice (indicated by different colors. **e**, Quantification of somata (LHb and off-target centrolateral thalamic nucleus) within the GRIN lens working distance (max. 250 μm) across mice. **f**, Mean calcium transients for all decisive trials. Smoothed data, 549 cells pooled across 12 mice, sorted for mean activity in the DP. Dashed lines indicate start and end of DP. **g**, Fractions of LHb neurons depending on their activity during the DP. **h**, Example traces from a positive (top) and negative (bottom) risk preference selective LHb neurons. **i**, Quantification of degree of selectivity (absolute values) for positive (n=196 cells) and negative (n=152 cells) risk preference selective cells (RPSCs). Correlation of DP responsiveness with DP selectivity for positive (blue) and negative (red) RPSCs for the preferred (**j**, positive: ρ=0.0137; p = 0.8486 and negative: ρ=0.2984; p = 0.00019) or non-preferred (**k**, positive: ρ= −0.6271; p < 0.0001 and negative: ρ=0.2984; p = 0.00019) choices. Correlation of DP activity difference between risky and safe choice (for positive (blue) and negative (red) RPSCs) with individual riskiness for present (**l**, positive: ρ=0.6294; p < 0.0001 and negative: ρ= −0.491; p < 0.0001) and previous choice activity (**m**, positive: ρ=0.2885; p = 0,74823and negative: ρ= 0.175; p = 0,0326).*p<0.05, **p<0.01, ***p < 0.001, Wilcoxon signed rank test (paired), χ^2^-test or Wilcoxon rank sum test (unpaired).

A potential confound for this interpretation might be differences in preparatory motor and licking behavior during the DP. Indeed we found that mice displayed more anticipatory licking for their preferred compared to their non-preferred option (Extended Data Fig. 8a). Thus we wondered whether this differences in motor activity might also influence LHb single cell activity. To test this, we correlated the DP activity of every recorded LHb neuron with the number of anticipatory licks during the DP on a trial-by-trial basis. Across mice we did not detect a correlation of LHb activity and anticipatory licking (Extended Data Fig. 8b-d). Next we extracted trials without anticipatory licks during the DP and isolated RPSCs from this reduced dataset (26.6±2.9 % of decisive trials). The relative fraction of positive and negative RPSCs, their selectivity, and the correlation of their mean DP activity with individual risk preference (Extended Data Fig. 8e-g), replicated our results from decisive trials including anticipatory lick trials (Fig. 3 g, i, l). Subsequently, we tested whether preparatory mouth movement activity would explain LHb single cell activity patterns during the DP. The correlation between mean LHb activity and mouth movement was elevated during the DP (compared to 1-s baseline before and 1-s post-deliberation period after the DP; Extended Data Fig. 8h). Furthermore, in line with augmented anticipatory licking for the individually preferred option, preparatory movement (as ratio of movement during the DP prior to risky and safe choices) was highly correlated to DP activity ratios (Extended Data Fig. 8i). However, mean LHb single cell activity did not align well with mouth movement and appeared prominently shifted with a lag of more than 0.5 s for about two-thirds of the cells (Extended Data Fig. 8j, k). Moreover, the differences in preparatory mouth movements for risky and safe choices did not correlate with individual risk preference (Extended Data Fig. 8l). In summary, while licking and preparatory movement reflect individual preference, they do not explain LHb single cell DP activity.

To test how positive and negative RPSCs would adapt once individual preference changes, we trained a subset of our GRIN implanted mice (n=4 mice) in a modified version of our task where we gradually altered the expected value ratio (Methods) between the animals’ non-preferred and preferred option, so that it became increasingly advantageous to shift to the alternative option, and tracked the activity of identified RPSCs over the course of choice adaptation (Extended Data Fig. 5a). Most RPSCs either inverted (43%) or lost (39%) their selectivity while a minority (18%) of RPSCs even increased their selectivity for the previously preferred option despite the animal having adapted its preference (Extended Data Fig. 9a,b). While the fraction of the latter population was equal for positive and negative RPSCs, the fraction of inverting RPSCs contained more positive than negative RPSCs, and the RPSCs which lost selectivity were almost exclusively positive RPSCs. Interestingly, while LHb RPSCs showed almost no selectivity during intermediate choice adaptation, selectivity was restored once the animals had established a new preference (Extended Data Fig. 9 c-h). These findings indicate that LHb RPSCs flexibly adapt to value-based changes and represent a preference which is habitual in nature.

### Divergent coupling of LHb and its input regions during risky decision-making

The LHb is known to receive direct synaptic input from multiple forebrain regions ^44, 45^ which contribute to distinct aspects of behavior. ^30, 31^ We hypothesized that excitatory synaptic input from some of these regions may modulate LHb selectivity during the DP in our risky decision-making paradigm. To evaluate anatomical connectivity of the LHb, we first employed retrograde adeno-associated virus (AAV) labelling of LHb inputs, whole-brain tissue clearing, and 3D light-sheet microscopy (Fig. 4a-e). Among the labeled areas were brain regions known to contribute to valence encoding and decision-making processes, ^17–20, 44, 46, 47^ such as the orbitofrontal cortex (OFC), areas of the medial prefrontal cortex (mPFC), the ventral pallidum (VP), as well as the lateral and medial hypothalamus (LH and MH). To verify that these regions indeed send excitatory axonal projections to the LHb, we injected anterograde viral labels into Vglut2-Cre mice. We found strong axonal labeling in the LHb from VP, LH, and MH (mainly originating from dorsomedial (DMH) but to a lesser extend also ventromedial hypothalamus (VMH), Supplementary Fig. 1b) injections, whereas the axonal projections from OFC and mPFC seem to rather target the adjacent centrolateral and mediodorsal thalamic nuclei (Fig. 4f-h; Supplementary Fig. 1).

**Fig. 4.**
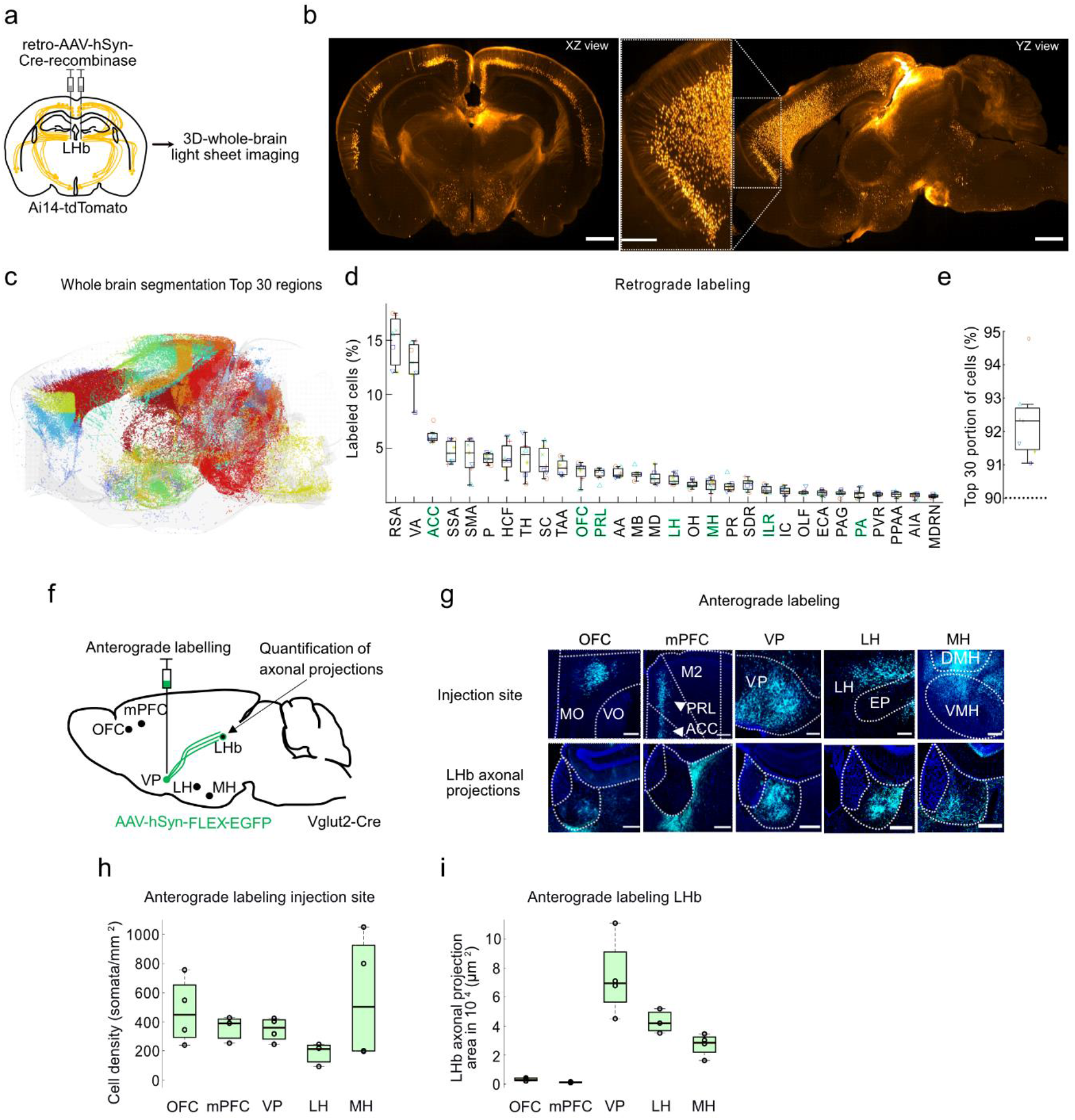
Retrograde and anterograde AAV-labelling of LHb inputs. **a**, Illustration of retrograde labelling approach.Example images (**b**) and segmented (**c**) retrogradely labelled LHb-projecting neurons across an entire mouse brain. Neurons are colored by brain areas; top 30 structures are shown here for the purpose of a clear visualization. Light grey outlines the brain surface. Different colors distinguish different brain regions. Scale bar represents 1 mm. **d**, Quantification of top 30 identified input regions (n=7 mice). Target areas for anterograde labelling and multi-fiber photometry highlighted in green. A detailed list of all abbreviations can be found in Extended Data Table 2. **e**, Coverage of cells detected in top 30 brain regions relative to all detected cells. **f**, Illustration of anterograde labelling approach. **g**, Examples of anterograde labelling of Vglut2-positive neurons and their axonal projections in the LHb from the five selected regions (orbital frontal cortex (OFC), medial prefrontal cortex (mPFC), ventral pallidum (VP), lateral hypothalamus (LH), medial hypothalamus (MH). Scale bars represent 200 μm. Quantification of glutamatergic somata (**h**) and axonal projection covered LHb area (**i**) arising from the five selected input regions (n=3-4 mice per region).

To examine how neuronal populations in the candidate input regions functionally couple to the LHb during our task, we next employed multi-fiber photometry implanting 2-6 fibers in Vglut2-Cre mice to simultaneously record task-related activity in the LHb and 1-2 input regions (n = 21 mice with 5-7 mice per region; Fig. 5 a-c; Supplementary Fig. 2). Based on our anatomical data, we expected VP, LH, and MH to couple most strongly to the LHb. Indeed, population activity in LH and MH yielded the highest correlation with LHb activity during the DP (Fig. 5d). VP showed some coupling to LHb, albeit weaker than for the hypothalamic inputs, whereas mPFC and OFC were rather uncoupled, consistent with the anterograde labeling results. Interestingly, whereas MH-LHb and LH-LHb coupling was similar for preferred and non-preferred choices, MH-LHb but not LH-LHb coupling increased significantly over the course of the DP (Fig. 5e, f), indicating potentially diverging functions of these two glutamatergic synaptic inputs. This aspect was further underlined by the divergent reward outcome encoding in MH and LH bulk activity. While both regions showed high inter-individual variability in reward responses (in the reward and post-reward period), MH bulk activity ratios for different reward outcomes generally showed higher correlation with risk preference behavior than LH bulk activity (Extended Data Fig. 10). To test whether MH and LH would resemble LHb population activity divergence during DP, we pooled multi-fiber recordings with data from animals implanted with a fiber in either MH or LH and compared the responses prior to preferred and non-preferred choices. While MH DP population activity did not show a significant difference (Fig. 5g, h; n=9 mice), LH DP population activity was significantly higher before the animals chose their preferred over the non-preferred option (Fig. 5g, i; n=9 mice). This divergence was particularly pronounced in the early DP (0 – 0.6 s) but to a lesser extend in the late DP (second 0.6 – 1 s; Fig. 3j, k). Similar to the LHb (Fig. 2f-h), LH DP activity ratio correlated with risk preference behavior (Fig. 5l), although to a lesser extent, indicating that the LHb does not solely relay LH information but rather integrates information from distinct long-range projections.

**Fig. 5.**
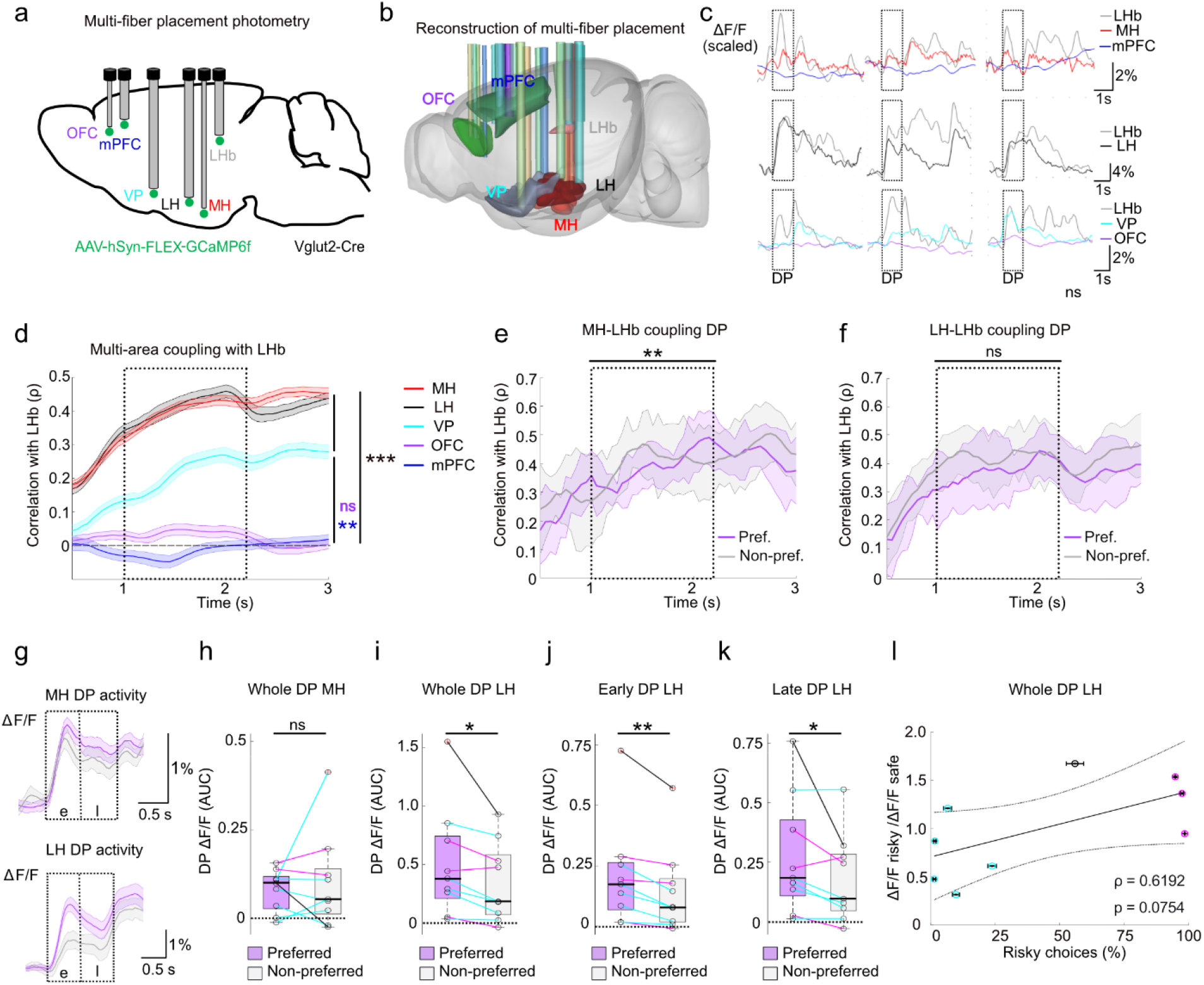
Functional coupling of long-range glutamatergic inputs and LHb is divergent. **a,** Illustration of multi-fiber-photometric recordings. **b**, Reconstruction of fiber placement across mice (n = 19 mice).Target areas are highlighted in different colors. **c**, Example of three single-trial multi-fiber-photometric calcium traces (columns) from three distinct animals (rows). Dashed boxes indicate DP. **d**, Coupling of LHb input region with LHb population activity (n=21 mice with 5-7 mice per region). Data represent coupling across sessions (291 in total) and include all trials where mice reported a response. Smoothed data are plotted as mean ± s.e.m. Coupling of MH (**e**) and LH (**f**) for preferred (violet) and non-preferred (grey) choice trials (n = 5 mice, each; compared is the effect of trial time on area-coupling over the DP). **g**, MH (top) and LH (bottom) population activity during the early (e) and late (l) DP prior to preferred (violet) and non-preferred (grey) choices (50 randomly selected trials for each condition, respectively). Data are plotted as mean ± s.e.m. **h**, Quantification of MH population activity integrals during the whole DP for preferred (violet) and non-preferred (grey) option (n = 9 mice). Quantification of LH population activity integrals during the whole (**i**), early (**j**), and late (**k**) DP (n = 9 mice). **l**, Ratio of LH ΔF/F integrals during the whole DP for present risky and safe choice population activity correlated with individual risk preference across sessions. *p<0.05, **p<0.01, ***p < 0.001, Two-way ANOVA and Wilcoxon signed rank test (paired).

### MH-projections to LHb control different aspects of risky decision-making

Next, we examined the behavioral relevance of glutamatergic LHb-projecting neuron activity in LH and MH during the DP. To selectively inhibit LHb-projecting populations, we bilaterally injected a retrograde AAV encoding Cre-dependent Flp-recombinase in the LHb of Vglut2-Cre mice and expressed a Flp-dependent variant of the proton-pump ArchT or EYFP in both hemispheres in either LH or MH (Fig. 6a-e; n = 8 mice for each region; n = 6 additional mice for each region with EYFP expression as controls). Subsequently, we randomly activated ArchT during the DP in 25% of trials and compared decision-making parameters for ArchT and EYFP-expressing mice within perturbation sessions (difference between light ON and OFF trials) to evaluate immediate trial-type specific effects and across sessions (difference between with and without light stimulation sessions; Fig. 6f; Methods) to evaluate perturbation effects on the general stability of performance over the course of the session. Compared to their respective EYFP control within perturbation sessions, light stimulation in ArchT expressing mice significantly decreased the animals’ decisiveness as well as their risk preference levels for inactivation of MH→LHb but not LH◊LHb neurons (Fig. 6g, h). Across sessions, decisiveness decreased in sessions with MH→LHb inhibition (Fig. 6i). Preference levels across sessions were not significantly altered, neither for LH→LHb nor for MH→LHb inhibition (Fig. 6j). Some individual animals that showed perturbation-induced alterations in across-session preference levels, turned out to have some degree of off-target labeling of glutamatergic entopeduncular nucleus (EP) neurons, which was correlated to performance impairment for this specific parameter (Fig. 6e; Suppl. Fig. 3). Taken together, these findings indicate that MH but not LH inputs convey information to the LHb prior to action selection required for risk preference-selective behavior in our task.

**Fig. 6.**
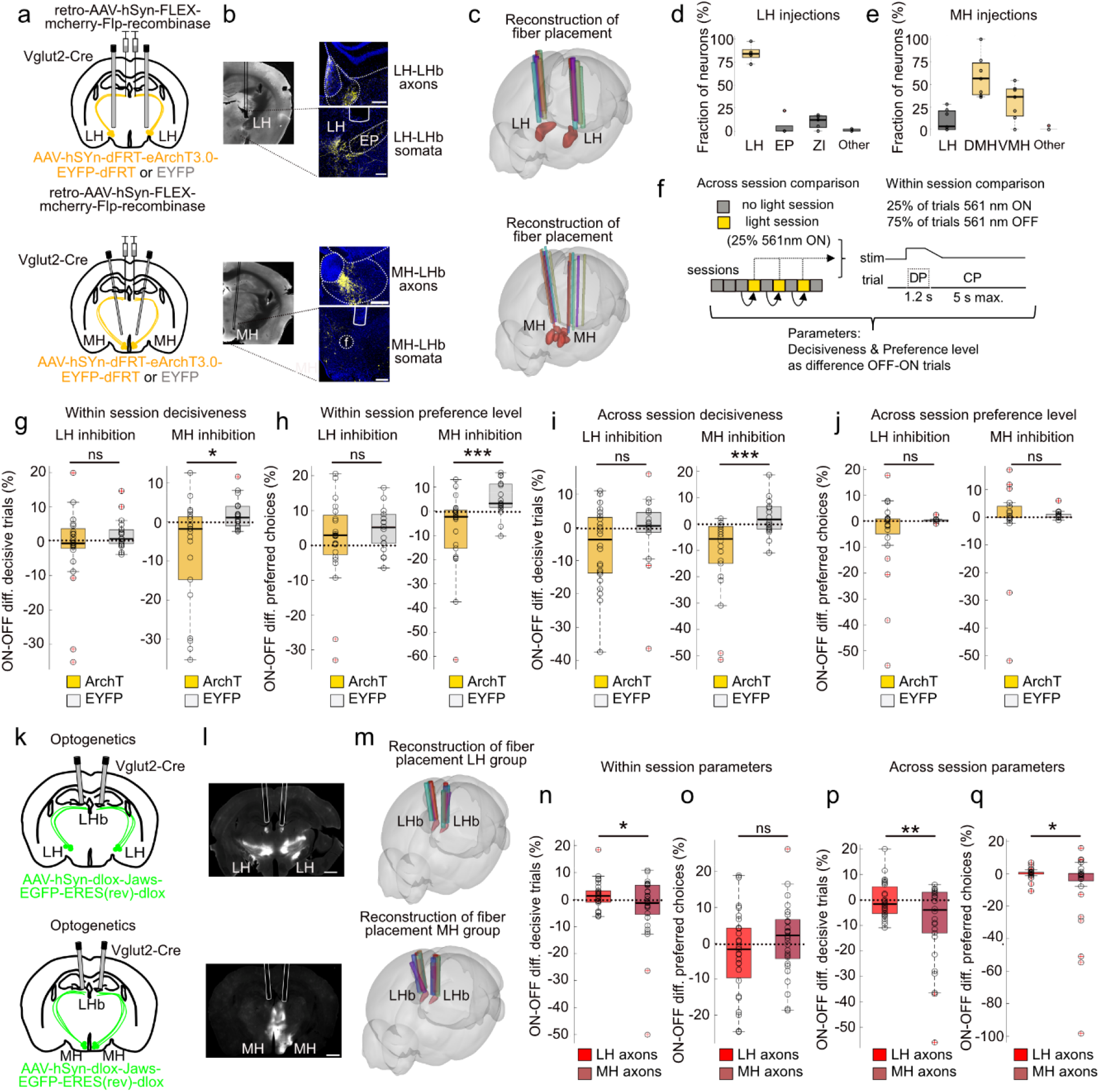
Functional distinct hypothalamus-LHb circuits are required for risky decision-making. **a**, Methodology for optogenetic inhibition of LH◊LHb and MH◊LHb projection neurons. **b**, Left: Widefield fluorescence images of half hemispheres, Right: Confocal images of retrogradely labelled ArchT-EYFP expressing neurons in the LH (top) and MH (bottom). Scale bars 200 μm. **c**, Reconstruction of fiber placement across ArchT-expressing mice (individuals indicated by different colors; n=7 for LH and MH). Quantification of labeled somata for LH (**d**) and MH (**e**) injected mice (entopeduncular nucleus (EP), zona incerta (ZI), dorsomedial hypothalamus (DMH), and ventromedial hypothalamus (VMH) across mice (n=5 for LH and n=7 for MH). **f**, Optogenetic perturbation protocol for across- and within-session comparisons. Labels indicate deliberation (DP) and choice period (CP). Within-session comparison of changes in decisiveness (**g**) and preference level (**h**) for ArchT-expressing mice (amber) and EYFP-expressing control mice (grey) for LH→LHb (left) and MH→LHb (right) projection neurons. **i**, Across-session comparison of changes in decisiveness for LH (left) and MH (right) inhibition. **j**, Across-session comparison of fraction of preferred choices for LH (left) and MH (right) inhibition (LH: 24 sessions from 8 ArchT mice, 18 sessions from 6 EYFP mice; MH: 22 sessions from 8 ArchT mice and 18 sessions from 6 EYFP mice). *p<0.05, ***p < 0.001, Wilcoxon rank sum test (unpaired). **k**, Illustration of LH◊LHb and MH◊LHb axonal perturbation approach. **l**, Widefield fluorescence images of hemispheres and reconstruction of fiber placement (**m**) targeting the LHb for inactivation of Jaws-expressing axonal terminals (individuals indicated by different colors; n=9 for LH and n=10 for MH). Within session comparison of changes in decisiveness (**n**) and preference level (**o**) for LH (left) and MH (right) axonal inhibition during DP. Across session comparison of changes in decisiveness (**p**) and preference level (**q**) for LH (left) and MH (right) axonal inhibition during DP (LH: 27 sessions from 9 mice; MH: 29 sessions from 10 mice). *p<0.05, **p < 0.01, Unpaired one-tailed t-test with Welch’s correction.

To test whether this information is indeed transmitted by MH projection neurons solely terminating in the LHb, we bilaterally injected a Cre-dependent version of the chloride-pump Jaws in either LH or MH and implanted optical fibers above the LHb to ensure LHb-specific axonal inhibition (Fig. 6 k-m, Methods). Direct comparison of LH◊LHb and MH◊LHb axonal inhibition during the DP revealed a robust impairment of decisiveness upon MH◊LHb axonal inhibition within and across sessions (Fig. 6 n, p). Moreover, we observed a subtle but significant effect of MH◊LHb axonal inhibition on across but not within session risk preference level stability (Fig. 6 o, q). In summary, retro- and anterograde perturbation experiments consistently show that MH◊LHb projections convey information which are required to execute confident choices during risky decision-making.

### LH and MH axonal projections to LHb show distinct functional properties

To better understand the differential behavioral effect of these two glutamatergic hypothalamic projections, we combined optogenetic inhibition of LH or MH axons in LHb with simultaneous recording of LHb population activity. Again, we bilaterally injected a Cre-dependent version of Jaws in either LH or MH and recorded LHb population calcium signals with fiber photometry with and without Jaws stimulation (Fig. 7a-c). For both projections, axonal inhibition by Jaws during the DP significantly reduced LHb calcium signals (Fig. 7d, e; decrement in ΔF/F integral: LH: − 0.08 ± 0.02 %·s, early, −0.19 ± 0.30 %·s, late; MH: −0.004 ± 0.025 %·s early, −0.08 ± 0.01 %·s late; median ± s.e.m.). However, whereas inactivation of LH axons rapidly induced a drop in LHb bulk fluorescence, the impact of MH axonal silencing was initially less prominent and occurred with a significant delay (Fig. 7f; onsets of drop: 0.18 ± 0.05 s in LH and 0.98 ± 0.24 s in MH, median ± s.e.m.). These results indicate that LHb activity depends differentially on synaptic inputs from MH vs. LH in the early vs. late phase of the DP.

**Fig. 7.**
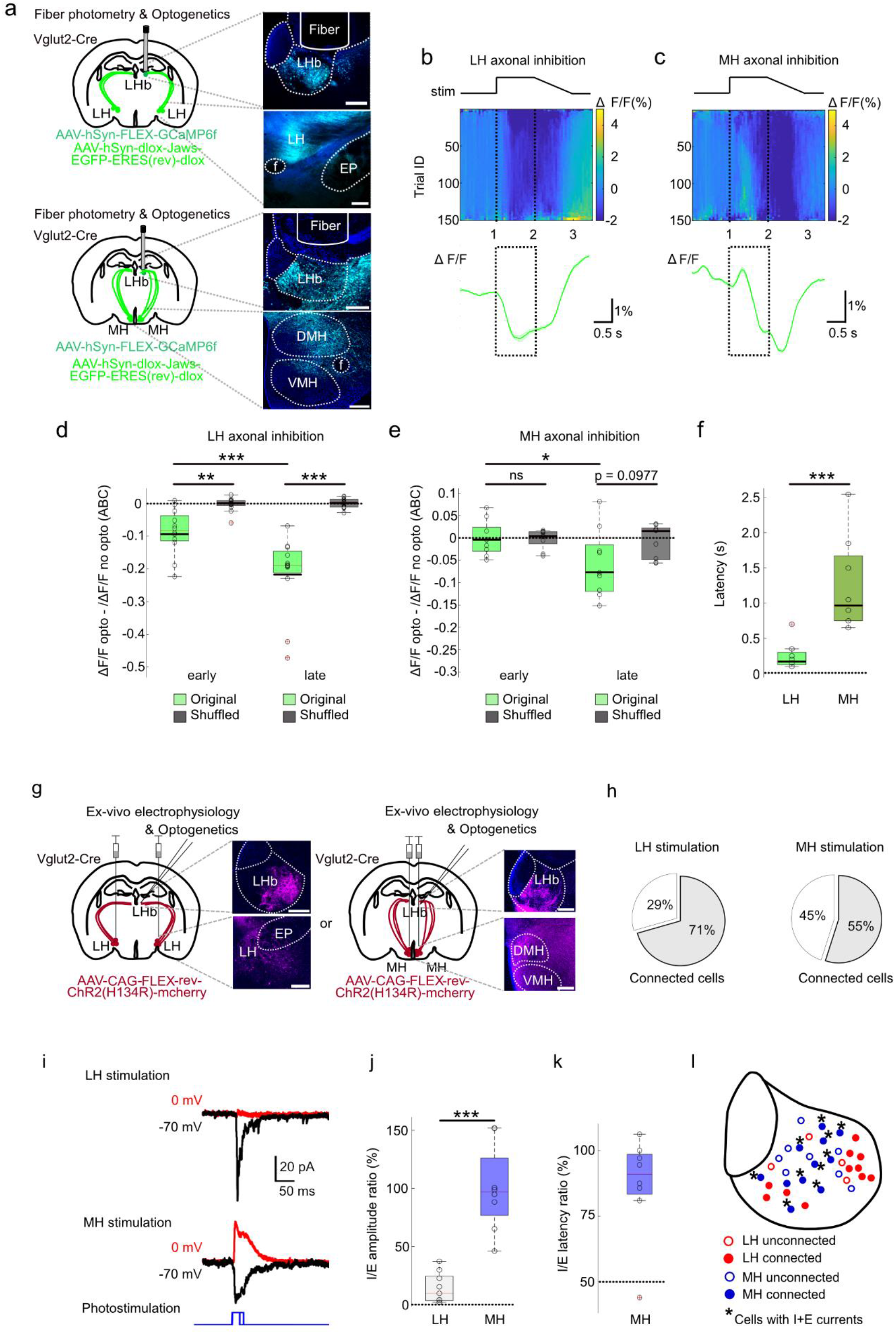
LH and MH axonal projections to LHb show distinct functional properties. **a**, Illustration of hybrid photometry/optogenetics experiment for either LH or MH axonal perturbation. Whereas LHb neurons express Cre-dependent GCaMP6f cytosolically, the Jaws-EGFP fusion protein is membrane-bound and thus leads to more diffuse labeling. **b**, Example LHb photometry difference traces (561 nm ON trace – 561 nm OFF trace) for LH (**b**) an MH (**c**) axonal inhibition. Difference between 150 randomly selected ON and OFF single trials (top; sampled across 3 sessions) and their respective mean (green trace; bottom; dashed box indicates DP.). Laser stimulation protocols shown above. Quantification of LHb population activity difference upon LH (**d**) and MH (**e**) axonal inhibition for early (0-0.6 s) and late (0.6-1.2 s) stimulation phase. Differences (with minus without inhibition) of photometric ΔF/F integrals in early and late window for original (green) and shuffled (grey) data. **f**, Latency to the onset of ΔF/F drop below pre-stimulation baseline for LH◊LHb and MH◊LHb axons (data from 12 sessions in 4 mice for LH and 9 sessions in 3 mice for MH). **g**, Illustration of ex vivo whole-cell recordings in brain slices with cell-type specific optogenetic stimulation of LH◊LHb or MH◊LHb axons. **h**, Fraction of LHb cells showing postsynaptic responses upon optogenetic stimulation of LH (12/17 cells from 4 mice) or MH (11/20 cells from 8 mice) axons. **i**, Example current traces of excitatory and inhibitory postsynaptic currents (EPSC, black, −70 mV holding potential; IPSC, red, 0 mV holding potential) evoked by photostimulation (blue) of LH→LHb (top) and MH→LHb (bottom) axons. **j**, Ratio of photostimulation-evoked IPSC/EPSC amplitudes (8 cells in 4 mice for MH axons; 9 cells in 4 mice for LH axons). **k**, Latency ratio for IPSC vs. EPSC evoked by MH axonal stimulation. **l**, Spatial mapping of LH (red) and MH (blue) connected (full circles) and unconnected (empty circles) recorded LHb neurons. Cells which showed inhibitory and excitatory responses are marked with an asterisk. *p<0.05, **p<0.01, ***p < 0.001, Wilcoxon signed rank test (paired) or Wilcoxon rank sum test (unpaired).

We next asked how Vglut2-positive LHb-projecting neurons in the LH and MH might exert such different influence on LHb neurons. To address this question, we expressed Cre-dependent channelrhodopsin-2 (ChR2) either in LH or MH of Vglut2-Cre mice and performed ex-vivo whole-cell recordings from LHb neurons in acute brain slices. We stimulated the ChR2-expressing axonal projections with light and measured the light-evoked synaptic currents in LHb neurons (Fig. 7g). Both LH◊LHb and MH◊LHb projections showed high connectivity with LHb neurons, as evidenced by light-evoked excitatory post-synaptic currents (EPSCs; Fig. 7h). Intriguingly, stimulation of MH◊LHb axons also elicited equivalent inhibitory post-synaptic currents (IPSCs) (Fig. 7i, j) whereas LH◊LHb axons showed GABA release only weakly and less frequently, as seen previously. ^44, 48^ MH-evoked EPSCs and IPSCs occurred with similar onset latencies (Fig. 7k; I/E latency ratio, 91 ± 62%), mainly in the medial part of the LHb (Fig. 7l), persisted under conditions of polysynaptic blockade (tetrodotoxin with 4-aminopyridine) and were sensitive to glutamate (Glu) receptor and GABA_A_ receptor antagonists (Suppl. Fig. 4). These findings suggest that Vglut2-positive MH◊LHb neurons co-release Glu and GABA in the LHb, whereas Vglut2-positive LH◊LHb neurons appear to release Glu principally ^44, 48^.

## DISCUSSION

Taken together, our study shows that evolutionary conserved brain regions form highly specific neuronal circuits to fulfill distinct functions in habitual risk preference-based decision-making. In particular, we uncovered that specific LHb neurons encode a risk preference decision bias prior to action selection. This bias is modulated by MH→LHb projection neurons which are capable of Glu and GABA co-release in the LHb and convey information about decisiveness but also preference.

### Probing risk preference in mice

We applied a balanced two-alternative choice task to test risk preference in head-fixed mice. Most animals displayed a strong risk preference which was stably maintained across weeks. The majority of individuals strongly preferred the safe over the risky option. Such prominent risk aversion has been observed in various other species, ranging from birds^1^ and rodents^2, 3^ over monkeys^4–6^ to humans^6, 7^. Mice further differed in their applied strategy to coping with loss. In line with previous work in rats^3^, risk averse mice showed higher loss sensitivity than risk prone mice, making them more likely to switch back to the safe option or show an indecisive choice and less likely to take the risky option again. Furthermore, like in rats^3, 10^, mice engaged in probabilistic discounting when expected value ratios between safe and risky option were altered reflecting flexible, value-based decision-making.

### Risk preference encoding in the LHb

In a previous study, pharmacological inactivation of the LHb impaired probabilistic discounting behavior by preventing the formation of value-based preferences^10^. However, how such value-based preferences are encoded in the LHb during risky decision-making remained elusive. Here, we found that LHb activity reflects individual risk preference during choice deliberation prior to action selection^41^, both at the population level and with cellular resolution. A subset of LHb neurons encoded individual risk preference even on the single-cell level, by displaying diverging activity for the individually preferred and non-preferred (be it risky or safe) choice options. In most neurons the activity was higher for the preferred choice option. Such ‘positive’ encoding of a preference is particularly remarkable given that increased LHb activity has largely been associated with negative valence signalling in aversive contexts^25, 32, 49, 50^. On the other hand, our findings are in line with other studies that found a subset of LHb neurons increasing their activity during reward anticipation^39, 43^.

### Hypothalamic modulation of LHb activity during risky decision-making

Furthermore, we identified two long-range hypothalamic projections providing excitatory inputs (and inhibitory for MH) that influenced LHb activity on the population as well as the single-cell level. However, in our task only MH→LHb projections which as we show here were capable of Glu and GABA co-release, bore behavioral relevance for risky decision-making. MH→LHb projections had been described anatomically ^51^ but were not functionally examined so far.

Glutamatergic LH→LHb projections were previously shown to encode negative motivational value and promote behavioral avoidance^44, 48^. Interestingly, a recent study found a specific subtype of Vglut2-positive, estrogen receptor 1-expressing LH→LHb projection neurons to be mainly responsible for driving behavioral avoidance^52^. In contrast to the MH, the LH sends glutamatergic and GABAergic inputs to the LHb via separate neuronal pools^44, 53^. This dichotomy is not limited to LH→LHb but extends to LH→VTA projections where GABAergic projections promote appetitive responses whereas glutamatergic projections drive behavioral avoidance^54, 55^. Here we found that while glutamatergic LH and LHb bulk activity was highly correlated and LH→LHb glutamatergic axonal perturbation potently impaired LHb bulk activity, perturbation of LH-LHb coupling during the DP did not yield behavior-relevant effects. One potential explanation for this might be that while glutamate release from LH axons evokes potent responses in LHb neurons, it is not sufficient to evoke RPSC selectivity. In contrary, Glu/GABA co-release from MH→LHb axon terminals might enable fine-tuned gain control^56^ over the activity of LHb RPSCs in particular during the late DP, setting a stable risk-preference bias to promote more confident decisions.

Across our anatomical and functional data we further found that LH→LHb axons tended to target the lateral, MH→LHb axonal projections preferentially target the medial LHb subdivision. Hence, we propose two parallel hypothalamic-habenula pathways preferentially targeting medial (MH) and lateral (LH) subdivisions of the LHb. Our proposed circuit model is further supported by the organization of LHb outputs. LHb projection neurons send direct, marginally overlapping axonal projections to dopaminergic and serotonergic neuromodulator systems in the ventral tegmental area (VTA), the medial raphe nucleus (MRN), and the dorsal raphe nucleus (DRN)^29, 31, 53, 57, 58^. Since these regions fulfill distinct functions in value-guided behaviors, we propose that the anatomically distinct hypothalamic-habenula inputs may contribute region-specific functions.

## Methods

All experimental procedures were carried out in accordance with the guidelines of the Federal Veterinary Office of Switzerland and were approved by the Cantonal Veterinary Office in Zurich under license number 156/2017, 234/2018, 153/2019, 213/2021, and 008/2022.

### Animals

A total of 144 male and 22 female (9 for axonal optogenetic inhibition and anxiety testing, 5 for axonal optogenetic excitation, 2 for multi-fiber photometry, and 6 for anterograde axonal labeling) adult mice were used in this study. Male and female mice where not trained at the same time but in separate batches. Mice were maintained on a reversed 12:12 light dark cycle and tested during the dark phase. For behavioral training we used C57BL6/J mice and Vglut2-ires-Cre mice (Jackson Laboratories, no. 016963). For fiber photometry, optogenetic perturbation, two-photon experiments, *ex vivo* electrophysiology, and anterograde tracing we used Vglut2-ires-Cre mice (133 mice). For *in vivo* electrophysiology and behavioral analysis we used C57BL6/J mice (26 mice). For tissue clearing we used ai14-tdtomato mice (Jackson Laboratories, no. 007908; 7 mice).

### Behavioral training

We trained animals in the balanced two-alternative choice task. On the first day of training, mice were shaped to sample both spouts using 2.5-µl sucrose rewards which were automatically delivered. Once the animals successfully learned to sample both spouts for reward, auto-reward was abandoned, and the spouts would deliver 2.5-µl rewards only upon licking after the reward cues. When mice learned this association and continued to sample both spouts, we would introduce free-choice sampling on day 2-3. Mice first performed 50 trials of forced choice, followed by 200-300 free choice trials where both spouts delivered the same reward quantity of 2.5 µl. After this initial training phase, different reward quantities were introduced to examine individual preferences. The ‘safe’ spout delivered a reward of 5 µl sucrose water in 100% of all cases, whereas the ‘risky’ spout either delivered 17 µl in 25% or 1 µl in 75% of cases. Hence the expected value was equal for the two waterspouts. Every mouse performed 50 forced choice trials where only one spout provided a reward with a maximum of 3 repetitions of the same option, followed by 200-300 free choice trials where the animal could freely decide which spout to sample and both spouts carried a potential reward. Start of the trial and end of the DP were signaled by a 6 kHz auditory tone (200ms each with a pause of 800 ms in between). Potential reward location was indicated by an LED light close to the respective spout, i.e., on one side in forced choice trials and on both sides for free choices. To exclude learning effects, mice were trained until their behavioral performance varied by less than 20% across more than three consecutive training days. Subsequently, every animal absolved additional 14 daily sessions consisting of 50 forced-choice and 200-300 free-choice trials. Mice required 2.33 ± 0.48 sessions (mean ± s.e.m., n = 52) before reaching stable performance levels. Moreover, mice were required to indicate a clear decision in more than 50% of all free choice trials (decisiveness > 50%). A trial was labelled indecisive when an animal sampled both spouts during the DP prior to action selection. Risk preference classification was determined based on the mice’s average engagement into risk across all 14 sessions (‘risk averse’, <25% risky choices; ‘risk neutral’, between 25%-75%; ‘risk prone’, >75%). These criteria were established by prior experimentation.

To test the influence of a potential site bias as well as the mice’s ability to measure expected value difference between spouts, we trained another 14 mice until they reached stable baseline performance and then inverted the site (left-right) of the risky and safe spout or altered the expected value of the safe spout (Fig. S1). Mice were trained until reaching a stable performance level before the expected value was further changed. During these site-bias and expected-value control experiments animals performed 50 forced-choice and 200 free-choice trials per daily session.

### Body movement measurements

Behavior was assessed in a sound-isolated box to shield the animals from external cues, except for two-photon imaging experiments (but with the microscope shielded from all sides). For assessment of general locomotor activity, we recorded the animals’ posture during the entire trial period with a frame rate of 30 Hz using an infra-red sensitive body camera (DMK23UP1300; The Imaging Source). To detect general body movements, we outlined three regions of interest covering the forelimbs, mouth, and neck area, respectively, as described previously ^59^. Next, we calculated body movement for each region and trial as 1 minus the frame-to-frame correlation as a function of time for each trial. For correlation with fluorescence data, data were down sampled to 20 Hz.

### Behavior analysis

To understand risk preference behavior in mice, we analyzed three main behavioral variables: Decisiveness, fraction of risky choices, and preference level. Decisiveness, *D*, was defined as the ratio of all trials, in which an animal clearly indicated a decision (i.e., ‘safe choice’, ‘risky choice’, or ‘miss’ trials), divided by the total number of trials, *T_tot_*:

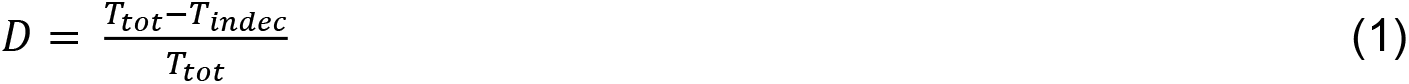

Where *T_indec_* denotes the number of indecisive trials. In the free-choice condition, a trial was labelled ‘indecisive’ when the animal sampled both spouts during the DP prior to action selection (and as a ‘miss’ if the animal did not lick at all). The fraction of risky choice, FRC, was defined as the ratio of all risky trials by the total number of all decisive trials:

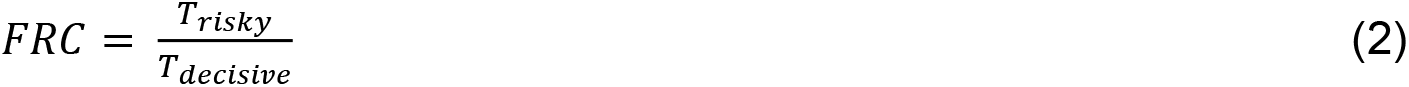

The preference level, PL, either for the safe or risky option was defined as the ratio of all preferred choice trials, *T_pref_*, by the total number of all decisive trials:

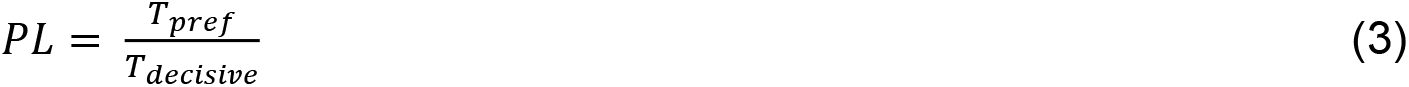

Behavioral modeling of choices was performed to better understand the history-dependence of risky or safe choices for each mouse. Modeling comprised four possible choices (risky choice, safe choice, indecisive, miss) and five possible outcomes (risky loss, risky win, safe, indecisive, miss). Transition matrices (Fig. S2B) describe the Markovian transition probabilities from the previous trial’s outcome to the current trial’s choice and were normalized such that the sum across all choices for a given outcome equaled to 1. To compare transition matrices across mice (Fig. S2E), the 4 x 5 transition matrix for each mouse was interpreted as a vector with 20 values and compared with transition matrices from other mice using Pearson’s correlation coefficient.

To investigate the history-dependence of choices for multiple preceding trials, we applied multinomial linear regression with the mnrfit() function in MATLAB, using the past outcomes to predict the current decision. The categorical variables used for regression were the possible decisions (‘safe choice’, ‘miss’, ‘indecisive’, ‘risky choice’), while the possible outcomes of the previous trials (‘safe choice’, ‘miss’, ‘indecisive’, ‘risky choice/loose’, ‘risky choice/win’) were used as regressors. To prevent overfitting, we used five-fold cross-validation. To quantify model performance, we computed a loss function that is used as standard for categorical variables (‘cross-entropy loss’) and computed the median performance across cross-validated subsets for each mouse. The loss function as a function of previous trials was normalized for each mouse by subtracting the minimum of the loss function. Plots in Fig. S2A show the negative of this normalized loss as the final performance metric.

### Open Field Test (OFT)

OFT testing took place inside a sound- and light-attenuating chamber (200 Lux; Omnitech Electronics, USA), retrofitted with a camera positioned above the chamber to enable video capture. Inside the chamber was a clear Perspex testing arena (41 x 41 x 30.5 cm). The testing arena contained soiled bedding material from all the mice that were tested and was not cleaned between trials (males and females were tested on different days and in soiled bedding from their own sex). All OFTs were 10 mins in duration.

### Light Dark Box (LDB)

LDB testing took place inside a LDB arena from TSE Systems Ltd, Germany. The LDB (internal dimensions: 42.5 cm (l), 29.5 cm (w), 24.5 cm (h) (dark compartment 15 cm (l) 29.5 cm (w) with a centered square opening 6 cm x 6 cm) consisted of both transparent and black infrared permeable Plexiglas walls and a light gray PVC floor. The LDB arena was placed on a table and recorded from above. Animals were tested under white light (100 Lux across the floor of the light compartment). An infrared light also illuminated the boxes so that an infrared camera could be used. Prior to testing each animal, the entire arena was cleaned using 70% ethanol. Animals were removed from their home cage and placed directly into the center of the light compartment. Tracking/recording was initiated once the animal was placed into the arena. All LDB tests were 10 mins in duration.

### Elevated plus maze (EPM)

EPM testing took place inside a custom built EPM. The EPM was made from black infrared permeable acrylic (walls (15cm high)) and matte white acrylic (floor), with arms measuring 65.5 × 5.5 cm (L × W), elevated 70 cm. Light intensity in the open arms was at 19–21 lux. All EPM tests were 5 mins in duration.

### Tracking and Analysis of OFT, LDB and EPM

The OFT, LDB and EPM were tracked using Deeplabcut and then analysed using the ETHZ DLC analyser^60^. Please see https://github.com/ETHZ-INS/DLCAnalyzer for more details.

### Virus injections

Before surgery, mice were briefly anesthetized with isoflurane (2%) in oxygen in an anesthesia chamber. Afterwards, animals were head-fixed in a stereotactic frame (Kopf Instruments) while anesthesia was maintained with 1-1.5% isoflurane. Body temperature was maintained at ∼37°C using a heating pad with a rectal thermal probe. To avoid dry eyes, the mouse’s eyes were covered with a vitamin A cream (Busch &Lomb). To prevent pain and inflammation, mice were subcutaneously injected with 0.1 μl/g body weight meloxicam. After longitudinal incision, connective tissue was removed, and the skull was dried using absorbent swabs (Sugi; Kettenbach).To target our regions of interest we used the following coordinates (from the brain surface): orbital frontal cortex (OFC, bregma (B): +2.80 mm, lateral (L): 0.2 mm, ventral (V): 1.2 mm), medial prefrontal cortex (mPFC, BLV: +1.70 mm, 0.2 mm, 1.2 mm), ventral pallidum (VP, BLV: +0.75 mm, 1.5 mm, 4.5 mm), medial hypothalamus (MH, BLV: −1.20 mm, 0.5 mm, 4.7 mm), lateral hypothalamus (LH, BLV: −1.30 mm, 1.6 mm, 3.6 mm), and lateral habenula (LHb, BLV: −1.70 mm, 0.5 mm, 2.5 mm). We used the following viral constructs and injection volumes per site in the different types of experiments: AAV.9.Syn.Flex.GCaMP6f.WPRE.SV40 (calcium recordings 140 nl; Addgene, no. 100833); AAV.1.hSyn.dio.EGFP (anterograde labelling, 140 nl, Addgene, no. 50457); AAV.9.hSy.chI.loxP.EGFP.loxP.SV40p(A) (anterograde labelling,140 nl, Viral Vector Facility of the University of Zurich (VVF), no. v56); retrograde.AAV.hSyn1.chI.mCherry.2A.iCre.WPRE.SV40p (retrograde labelling of LHb inputs and whole-brain clearing, 70 nl, VVF, no. v147), retrograde.AAV.hSyn1.chI.dlox.mCherry.2A.NLS.FLPo(rev).dlox.WPRE.SV40p(A) (cell-type specific retrograde labelling, 140 nl, VVF, no. v173), ssAAV1.2.hSyn1.chI.dFRT.eArchT3.0.EYFP.dFRT.WPRE.hGHp(A) (optogenetics, 140 nl, VVF, no. v423), ssAAV.5.2.hSyn1.dlox.Jaws.KGC.EGFP.ERES(rev).dlox.WPRE.bGHp(A).SV40p(A) (optogenetics, 140 nl, VVF, no. v508), AAV.9.FLEX.rev.ChR2(H134R).mCherry (optogenetics, 140 nl, Addgene, no. 18916), AAV.9.Ef1A.fDIO.EYFP (control for optogenetics, 140 nl, Addgene, no55641).

### Fiber and GRIN lens implantation

Upon removal of skin and connective tissue, iBond (Kulzer, Total Etch) was applied to the clean skull and cured by UV light. Subsequently, a slim ring of Charisma (Kulzer, A1) was sculpted, and UV cured at the edge of the exposed skull to optimize implant stability. Small craniotomies were drilled above the regions of interest. Fiber implants or 600-µm gradient index (GRIN) lens (Inscopix) were slowly lowered right above the respective regions using custom-made holders and fixed in place using UV cured dental cement. Fiber implants consisted of an optical fiber (200-µm or 400-µm core) attached to a metal ferrule (Thorlabs).

### Fiber photometry

Fiber photometry recordings were performed with a custom-built setup that allowed simultaneous recording of three channels in parallel. For excitation we used an Omicron LuxX 473-nm laser, modulated at 490 Hz. Laser light was attenuated with a neutral density filter (NDC-25C-4M, Thorlabs) and split twice using non-polarizing beam splitters (CCM1-BS013/M, Thorlabs). For detection we used photomultiplier tubes (Hamamatsu). Fluorescence emission was recorded at 2 kHz and downsampled to 20 Hz for analysis. Excitation and emission light were transmitted through the same multimode fiber (UM22-200 and UM22-400, NA = 0.22; Thorlabs). For combination of fiber photometric recordings with optogenetic perturbation, 561-nm laser light (Coherent, OBIS) was coupled into the light path of one channel.

### Optogenetics

For optogenetic perturbation, we used a Cre-dependent version of the chloride pump Jaws as well as a Flp-dependent version of the proton pump ArchT. To assess stable baseline performance prior to optogenetic intervention, multiple baseline sessions without opsin activation were recorded. For ArchT perturbation and Jaws-photometry hybrid experiments, opsins were activated using 561-nm laser light (OBIS laser; Coherent) for Jaws perturbation experiments 633-nm laser light (OBIS laser; Coherent) was used. To circumvent excessive tissue heating ^61^ as well as rebound activation ^36, 62^, low intensity laser light (5-6 mW at the fiber tip) was modulated at 40 Hz and ramped down for 1 s. Moreover, perturbation was randomly applied in a maximum of 25% of trials within a session. Every perturbation session was followed by a session without opsin activation (non-perturbation session).

We compared decision confidence and preference level within a perturbation session as well as decision confidence and preference level stability between perturbation and a non-perturbation session. We evaluated the following changes for light ON compared to light OFF trials:

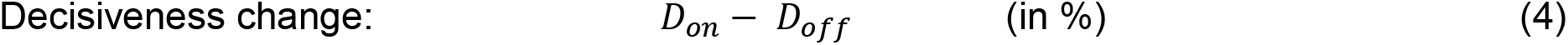

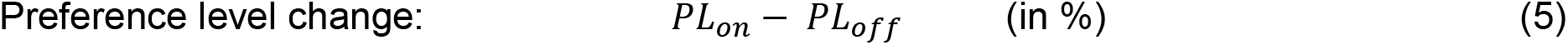

These changes were compared between EYFP-expressing and ArchT-expressing mice.

### *In vivo* electrophysiology

Chronic electrophysiological recordings were performed in five wild-type (C57BL/6) male mice. During the initial surgery, the scalp was retracted, the skull was exposed and the implantation target was marked with a permanent marker before sealing the skull with dental acrylic and positioning a headpost over the cerebellum. Mice were subsequently allowed to recover for 7 days before they were handled and acclimatized to the behavioral setup and trained on the task. In a second surgery, the mice were anesthetized as previously and a silicon linear electrode array (16 sites, 20-μm inter-electrode distance; Atlas Neurotechnologies) was targeted to the LHb. The linear array was dipped in a lipophilic dye (DiI; Sigma-Aldrich) for subsequent localization. A small craniotomy (∼0.75 mm) was performed above the target and two additional trepanations (∼0.2 mm) were performed over the cerebellum and the frontal cortex for placement of ground and reference electrodes. Next, the probe was slowly inserted centering the electrode sites in the LHb. After implantation the probe was fixed in place with additional acrylic and silver wires placed in contact with the cerebrospinal fluid (CSF) to serve as ground and reference electrodes. The mice were allowed to recover for another 5 days after surgery and their health was monitored. Mice were placed in an enclosed, sound-proof box and head-fixed during performance of the task. Electrophysiological recordings were performed while the mice performed the task. The voltage from the electrode array was amplified and digitally sampled at a rate of 30 kHz using a commercial extracellular recording system (Intan). The raw voltage traces were filtered off-line to separate the multiunit activity (MUA) (band-pass filter 0.6–6 kHz) using a fifth-order Butterworth filter. Subsequently, for each electrode a threshold was applied to the high-pass filtered data to isolate multiunit activity and reject background noise (4 times the standard deviation across the recording session, specified per electrode). The MUA was subsequently downsampled to the sampling rate of the fiber photometry (20 Hz) to compare the population time course with the bulk photometry signal.

For isolation of single units, the raw voltage traces were filtered off-line to separate the MUA (highpass filter frequency of 480 Hz) using a second-order Butterworth filter. Subsequently, for each electrode, a threshold (4x the STD a given electrode) was applied to the highpass filtered data to isolate single spike events and the resulting waveforms in a 2ms window around the threshold crossing where used to identify putative single units. Single units were distinguished based on their amplitude and waveform shape using the JRCLUST toolbox^63^ (https://www.biorxiv.org/content/10.1101/101030v2). The firing rates for single units isolated for individual mice where then sorted based on present or previous trial choice and compared with appropriate statistics during the DP.

### *Ex-vivo* electrophysiology

Whole-cell patch-clamp electrophysiological recordings were obtained in LHb-containing acute brain slices prepared from adult mice. Coronal brain slices were prepared in cold choline cutting solution containing (in mM): 100 choline chloride, 25 NaHCO_3_, 25 D-glucose, 2.5 KCl, 7 MgCl_2_, 0.5 CaCl_2_ and 1.25 NaH_2_PO_4_, aerated with 95% O_2_ / 5% CO_2_. Slices were perfused at a rate of 0.5–1 ml/min with oxygenated artificial CSF (aCSF) containing (in mM): 128 NaCl, 26 NaHCO_3_, 10 D-glucose, 3 KCl, 1 MgCl_2_, 2 CaCl_2_ and 1.25 NaH_2_PO_4_, aerated with 95% O_2_ / 5% CO_2_ at 29°C. Patch electrodes were made from borosilicate glass (Harvard Apparatus) and had a resistance of 2-4 MΩ. The intracellular solution contained (in mM): 126 cesium methanesulfonate, 4 CsCl, 10 HEPES, 20 phosphocreatine, 4 MgATP, 0.3 NaGTP, pH 7.3, 290 mOsm. Experiments were performed in voltage-clamp mode using the Axopatch 200B amplifier (Molecular Devices). Visually-guided whole-cell recordings were obtained of cells in a field of fluorescent axons (from AAV.9.FLEX.rev.ChR2 (H134R).mCherry injection into LH or MH of Vglut2-cre animals) on a Zeiss Axioscope using an optiMOS camera (Q Imaging). Access resistance was monitored to ensure the stability of recording conditions. Recordings with access resistance >40 MΩ or whole cell capacitance <4 pF were excluded. No compensation was made for access resistance and no correction was made for the junction potential between the pipette and the ACSF. Following a baseline stabilization period (2-3 min), electrically isolated excitatory or inhibitory postsynaptic currents were evoked by photostimulation (488 nm) at holding membrane potentials of V_h_ = −70 mV and V_h_ = 0 mV, respectively. Nine recurring photostimuli of sequentially increasing length (20 to 250 ms) were delivered at 0.2 Hz with a Polygon400 (Mightex), localized to the soma of the recorded cells. An average of 30 sweeps were analyzed for each condition from each cell using Clampfit10 (Molecular Devices). Bath applied compounds TTX (1 µM, Tocris 1069, CAS: 18660-81-6), 4-AP (100 µM, Tocris, 0940, CAS: 504-24-5), CNQX (20 µM, Tocris, 1045, CAS: 479347-85-8), D-APV (100 µM, Tocris, 0106, CAS: 79055-68-8), or gabazine (25 µM, Tocris, 1262, CAS: 104104-50-9) were dissolved as stock solutions in water. For quantification, peak current amplitude was calculated as an average of 3 ms surrounding the first evoked peak. Onset delay was measured as the average time to initial deflection from stimulus onset. Average spiking was calculated as the average spike rate of 2x 20-ms windows prior to (pre-stim.) and during (stim.) optogenetic stimulation.

### Two-photon calcium imaging

Two-photon calcium imaging data were acquired with a custom-built two-photon microscope controlled by ScanImage (Basic Version; Vidrio Technologies LLC, Ashburn), equipped with a Ti:sapphire laser system (approximately 100-fs laser pulses, center wavelength 910 nm; Mai Tai HP, Newport Spectra Physics), a Pockels cell for intensity modulation (Conoptics), a laser scanning unit with an 8-kHz resonant scan mirror (Cambridge Technology) and a galvanometric mirror (6215H, Cambridge Technology), and a 20x air objective (N Plan L 566035, NA = 0.40; Leica). Imaging was performed through the implanted GRIN lens (500-µm diameter, ∼6.1 mm length, 1050-004599, Inscopix). Fluorescence signal was detected with a photomultiplier tube (Hamamatsu) after passing through an IR blocking filter. Images were acquired at 30 Hz with 512x512 pixel resolution. Laser power was adjusted to 30 mW under the objective. Single trials (8.2 – 12.7 s long) were recorded with breaks of 0.5 s during the inter-trial period to allow data saving. To ensure two-photon signal integrity during the two-alternative choice task, LED signalling was omitted in these experiments and the reward cue solely consisted of the 6-kHz auditory tone.

### Source extraction and quality control of calcium imaging data

Raw data were corrected for brain motion using a modified version of rigid movement correction based on phase correlation (Suite2p)^64^. Under the GRIN lens, the illumination profile is highly inhomogeneous across the field of view (FOV), with higher intensity in the center of the FOV and lower intensity at the edges. To avoid brain motion artifacts due to the illumination profile that moved by the same amount, we estimated the illumination profile by smoothing the average fluorescence image with a 2D median filter (100-pixel size). We divided each raw movie frame by this illumination template, performed rigid movement correction, and then multiplied each corrected movie frame with the illumination template. For source extraction, a trial-averaged movie was generated for each session and used to manually draw regions of interest (ROIs) around active neurons ^65^. Neuronal ROIs were manually mapped across session based on visual inspection of the mean fluorescence image, and discarded if the respective ROI had moved out of focus for a new session. ROIs were drawn based on anatomical features visible in the average fluorescence image, but also based on maps of local correlations ^66^. Potential residual movement artifacts were identified by inspection of the map of local correlations. Movement artifacts were usually visible as anti-correlated left/right or top/bottom borders of a single neuron. To rule out motion artifacts from a second perspective, trial-averaged movies and single-trial movies were visually inspected together with the extracted traces. Mice for which movement artifacts could not be eliminated, e.g. due to excessive axial brain motion, were excluded from the analysis.

### Calcium imaging data analysis and functional classification

Calcium transient time series data were extracted and analyzed using custom MATLAB code. ΔF/F was calculated on a trial-by-trial basis for each ROI as: (F - F_0_)/F_0_, where baseline F_0_ is the average signal over the last second of the inter-trial interval before the next trial starts with the auditory starting cue. To identify responsive neurons that show a substantial increase or decrease in fluorescence signal during the DP, we compared the mean calcium activity of the last second of the inter-trial interval with the mean calcium activity of the DP (z-scored data) of all trials where mice indicated a clear decision. Cells whose mean calcium activity was at least ten times higher or lower in the DP than in the baseline period classified as responsive cells.

To identify the ‘risk-preference-selective cells’ (RPSCs) amongst all responsive cells, we first calculated the area under the curve (AUC) for z-scored ΔF/F traces during the DP for preferred and non-preferred choice trials. We then drew a bootstrapped data sample and calculated a bootstrap statistic for each trial type using MATLAB ‘bootstrp’ function. Subsequently, we calculated the difference between randomly selected preferred and non-preferred trial DP activity and statistically compared them (Wilcoxon signed rank test, p<0.05) to the difference calculated from the same data sample with shuffled trial labels. Finally, we calculated a receiver-operating characteristic (ROC) curve for all significantly diverging cells during the DP. All cells with an AUC value >0.65 were considered to be RPSCs.

To evaluate the activity of RPSCs upon preference adaptation, we changed the expected value (EV) ratio between the two options by gradually decreasing the value of the preferred and increasing the value of the non-preferred option, tracked RPSCs identified in five baseline session prior to EV adjustment and calculated their selectivity during the DP as described above. Only cells, for which could be tracked throughout the entire period were included in the final analysis.

### *Post hoc* immunohistochemistry, probe location, and assessment of cellular distribution

After completion of behavioral training, mice were anesthetized (100 mg/kg and 20 mg/kg xylazine) and perfused transcardially with 4% paraformaldehyde in phosphate buffer, pH 7.4. After perfusion, tissue was removed from the ventral side of the skull bone until the brain was exposed. Subsequently, the exposed brain with the respective implant was additionally post-fixed with 4% paraformaldehyde for 2 to 7 days. After fixation, the brain was carefully separated from the implant. Coronal sections (100 µm thick) were cut with a vibratome (VT100; Leica). Sections were mounted onto glass slides and confocal z-stacks or widefield images were acquired using an Olympus FV1000 or a Zeiss Axio Scan Z1, respectively. For validation of probe location we aligned widefield images of hemispheres to the Allen Mouse Brain Common Coordinate Framework (version 3, 2017)^67^ using the QuickNII tool^68^ on the EBRAINS Collaboratory infrastructure. Subsequently we annotated probe locations and visualized the results using brainrender (BrainGlobe) ^69^ and custom-written Python code. To assess the spatial distribution of cells contributing to respective signals or behavioral effects, we annotated cells in confocal z-stacks below the determined center of the probe tip using the Fiji (Image J, 1.8.0_172 64 bit) plugin cellcounter. Based on anatomical landmarks, we annotated cells up to the maximum depth to which they could theoretically contribute to a given signal (200 µm for fibers with 200 µm core, 400 µm for 400 µm core fibers^70, 71^, and 250 µm for GRIN lenses (Inscopix)).

### Anterograde tracing data acquisition and analysis

After three weeks of viral expression, mice were sacrificed, the brain was extracted, and coronal sections were prepared as described above. Subsequently, we performed confocal imaging acquiring z-stacks of the respective injection site to quantify somatic labelling or the LHb to detect axonal projections from putative input regions, using the 10x objective of the Olympus FV1000. Data were post-processed and quantified by using Fiji (Image J, 1.8.0_172 64 bit).

### Tissue clearing, light-sheet imaging and data analysis

Brain samples were stained for tdTomato and cleared with a modified version of the iDISCO protocol ^72^. After six weeks of expression, mice were perfused and the brains post-fixed in 4% PFA in PBS for 4.5 hours at 4°C, shaking at 40 rpm. Brains were washed in PBS for three days at RT and 40 rpm, with daily solution exchange. Samples were dehydrated in serial incubations of 20%, 40%, 60%, 80% methanol (MeOH) in ddH_2_O, followed by two times 100% MeOH, each for one hour at RT and 40 rpm. Pre-clearing was performed in 33% MeOH in dichloromethane (DCM) overnight (o.n.) at RT and 40 rpm. After two times washing in 100% MeOH each for 1 hour at RT and then 4°C at 40 rpm, bleaching was performed in 5% hydrogen peroxide in MeOH for 20 hours at 4°C and 40 rpm. Samples were rehydrated in serial incubations of 80%, 60%, 40%, and 20% MeOH in in ddH_2_O, followed by PBS, each for one hour at RT and 40 rpm. Permeabilization was performed by incubating the mouse brains two times in 0.2% TritonX-100 in PBS each for 1 hour at RT and 40 rpm, followed by incubation in 0.2% TritonX-100 + 10% dimethyl sulfoxide (DMSO) + 2.3% glycine + 0.1% sodium azide (NaN_3_) in PBS for three days at 37°C and 65 rpm. Blocking was performed in 0.2% Tween-20 + 0.1% heparine (10 mg/ml) + 5% DMSO + 6% donkey serum in PBS for two days at 37°C and 65 rpm. Samples were stained gradually with primary polyclonal rabbit-anti-RFP antibody (Rockland, 600-401-379-RTU) and secondary donkey-anti-rabbit-Cy3 antibody (Jackson ImmunoResearch, 711-165-152) 1:2000 in 0.2% Tween-20 + 0.1% heparine + 5% DMSO + 0.1% NaN_3_ in PBS (staining buffer) in a total volume of 1.5 ml per sample every week for four weeks at 37°C and 65 rpm. Washing steps were performed in staining buffer five times each for one hour, and then for two days at RT and 40 rpm. Clearing was started by dehydrating the samples in serial MeOH incubations as described above. Delipidation was performed in 33% MeOH in DCM o.n. at RT and 40 rpm, followed by two times 100% DCM each for 30 minutes at RT and 40 rpm. Refractive index (RI) matching was achieved in dibenzyl ether (DBE, RI = 1.56) for four hours at RT. 3D stacks of cleared brains and hemispheres were acquired using the mesoSPIM light-sheet microscope^73^ at 1X with a resolution of 6 µm/voxel. Imaging data were post-processed by using Fiji (Image J, 1.8.0_172 64 bit) and Imaris (9.8.0, Oxford Instruments), and analyzed by using ClearMap2^74^ (https://github.com/ChristophKirst). The analysis consisted of three steps: alignment to standard atlas, cell detection, and segmentation into annotated brain areas. Briefly, the autofluorescence channel of cleared brain samples was first aligned to the standard Allen brain atlas (http://www.brain-map.org). Then, potential cells were detected by finding local maxima in the 3D volume. Detected cells were filtered by size with a custom threshold adjusted to each sample. Finally, detected cells were assigned to brain regions based on the annotated brain atlas. All brain samples were manually screened by two experimentalists to ensure the quality of alignment and cell detection, and spurious cell detection from structures with no clear cell labelling is excluded from final analysis.

### Statistical analysis

Statistical analyses are described in the main text, Fig. legends, and Extended Data Table 2. We applied parametrical (two-sided t-test) or non-parametrical (Wilcoxon rank sum test, Wilcoxon signed rank test, Chi square test) tests in MATLAB 2019a and two-way ANOVA, Friedman-test, and unpaired t-test with Welch’s correction in GraphPad Prism 5. To account for unequal sample size when comparing preferred vs. non-preferred signals, we drew equal sized bootstrapped data samples and calculated a bootstrap statistic for each trial type using MATLAB ‘bootstrp’ function. Significance was assumed as p<0.05.

## Data and code availability

Data and code will be made publicly available upon publication.

## Contributions

D.G. conceived the project; D.G. and F.H. designed the study; D.G. carried out all surgeries, behavioral training, and in vivo experiments with the help of Y.S. (fiber photometry), C.L. (multi-unit electrophysiology), and P.R. (two-photon imaging); D.G.,A.R., O.S.,P.R., and C.L. analyzed behavioral, optical, and *in vivo* electrophysiological data; A.M.R. performed tissue clearing and light-sheet imaging together with D.G.; S.H. and D.G. analyzed light-sheet data; T.S. performed and analyzed ex vivo brain slice recordings; D.G. and M.W. designed, and M.W. wrote the custom behavior control software; J.B., T.K., A.A., and F.H. provided supervision; D.G. and F.H. wrote the manuscript with comments from all authors.

## Acknowledgements

This work was support by grants to F.H. from the Swiss National Science Foundation (project grants no. 310030-127091 and 310030_192617; Sinergia grant no. CRSII5-18O316), the European Research Council (ERC Advanced Grant, BRAINCOMPATH, 670757), and to D.G., C.L., P.R., and S.H. from the University of Zurich (Forschungskredit grants, projects K-41220-05-01, K-41220-04, K-41220-06-01, and K-41220-07-01).). F.H. and T.K. received funding from the University Research Priority Program (URPP) ‘Adaptive Brain Circuits in Development and Learning’ (AdaBD). The authors thank Philipp Bethge for managing transgenic mouse lines, Dubravka Dujmovic-Göckeritz for genotyping, the 3R Hub at the ETH Zurich for their support with the additional behavior experiments and analysis, Hansjörg Kasper and Stefan Giger for technical support, Christian Ruff, Philippe Tobler and Manuel Mameli for helpful discussions.

## Extended Data Figures and Tables

**Extended Data Fig. 1.**
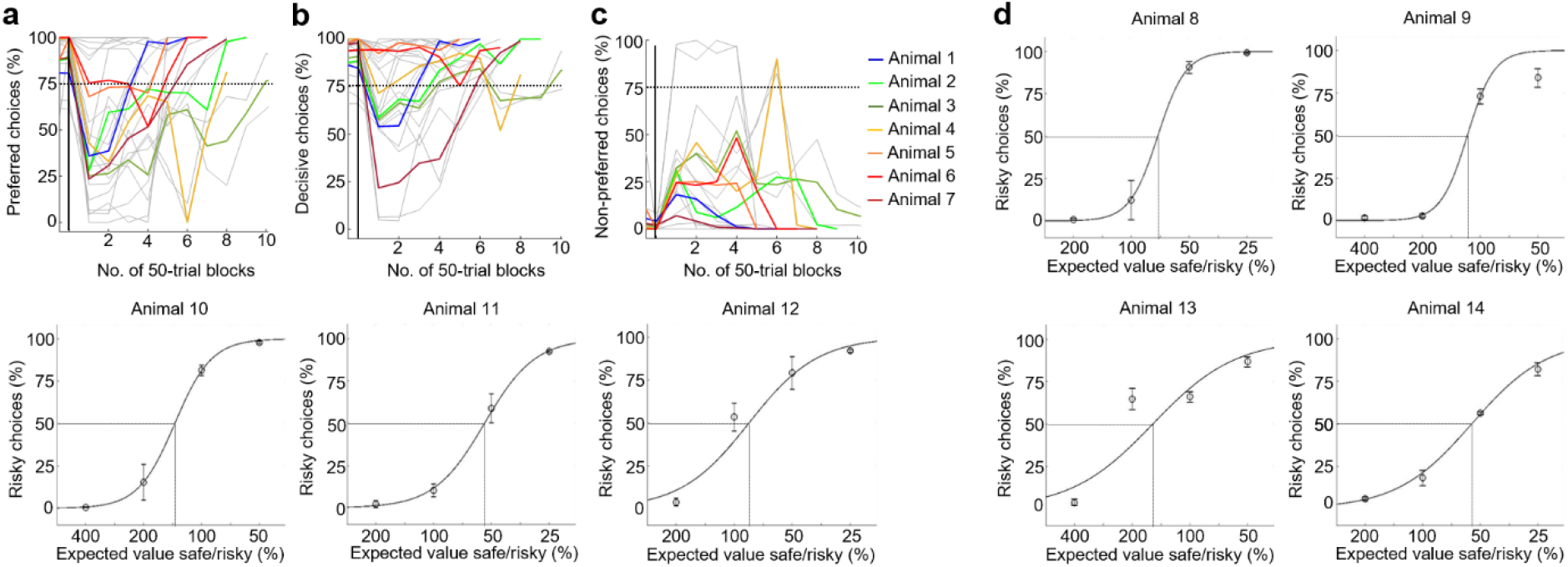
Mice performing two alternative choice task adapt their choices depending on preferred spout orientation and expected values. Fraction of preferred (**a**), decisive (**b**), and non-preferred (**c**) choices aligned to spout inversion (solid vertical line). Grey lines indicate individual inversion sessions (2-4 spout inversions per mouse), colored lines indicate mean across sessions for individual animals, dashed line indicates preference threshold. **d,** Every plot represents data from an individual mouse in response to decreasing value of the safe option. Every black dot represents the mean risky choices across four stable sessions with 200 trials each. Dotted lines indicate indifference point for each individual. Sigmoid fit.

**Extended Data Fig. 2.**
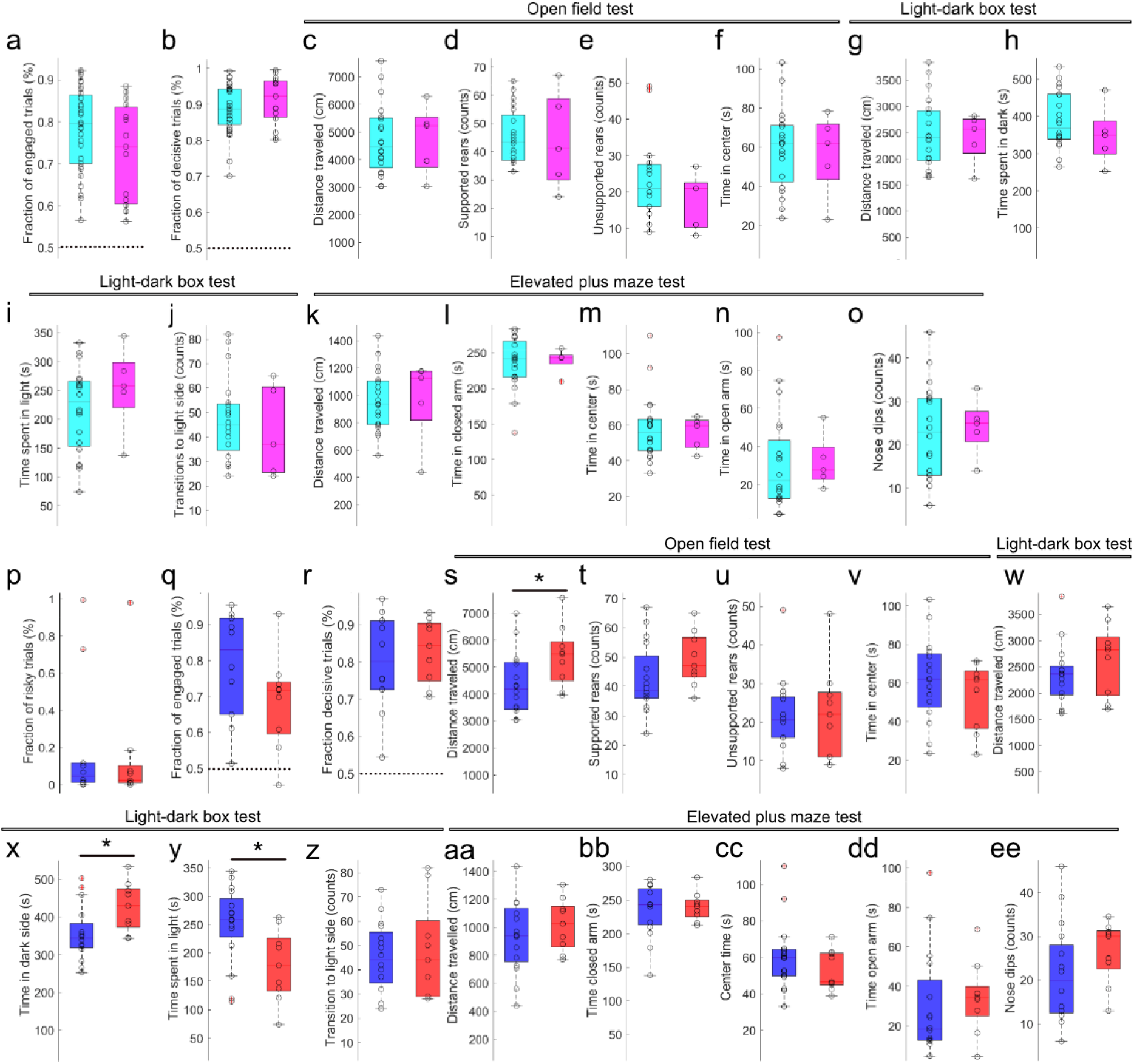
Risk preference is independent of motivation, decisiveness, anxiety or sex. **a**, Fraction of engaged trials for all risk averse (cyan, n=32) and risk prone (magenta, n=15) mice. Open circles indicate fractions of engaged trials (decisive and indecisive) from all trials an animal could engaged in (200 or 300 per session in 14 sessions, 2800 or 4200 trials for each mouse; p=0.1076). **b**, Fraction of decisive trials measured as the fraction of all trials an animal could engage in subtracted by the number of indecisive trials (p=0.2178). Results from plus maze test of animals after completion of two-alternative choice task. Comparison of risk averse (cyan, n=21) and risk prone (magenta, n=4) mice in open field (**c**-**f**; p=0.9188, p=0.7596, p=0.3948, p=0.9729), light-dark (**g**-**j**; p=0.9729, p=0.3958, p=0.3591, p=0.5633), and elevated plus maze test (**k**-**o**; p=0.4756, p=0.9188, p=1.0, p=0.5187, p=0.7084). Fraction risky (**p**, p=0.6207), engaged (**q**, p=0.1564), and decisive (**r**, p=0.8421) trials for male (blue, n=10) and female (red, n=9) mice (based on 600 trials across 3 sessions). Comparison of the same animals in open field (**s**-**v**; p=0.0338, p=0.1063, p=0.8872, p=0.4117), light-dark (**w**-**z**; p=0.1834, p=0.0253, p=0.0219, p=0.8650), and elevated plus maze test (**aa**-**ee**; p=0.4791, p=1.0, p=0.3802, p=0.3502, p=0.1484).*p<0.05, Wilcoxon rank sum test.

**Extended Data Fig. 3.**
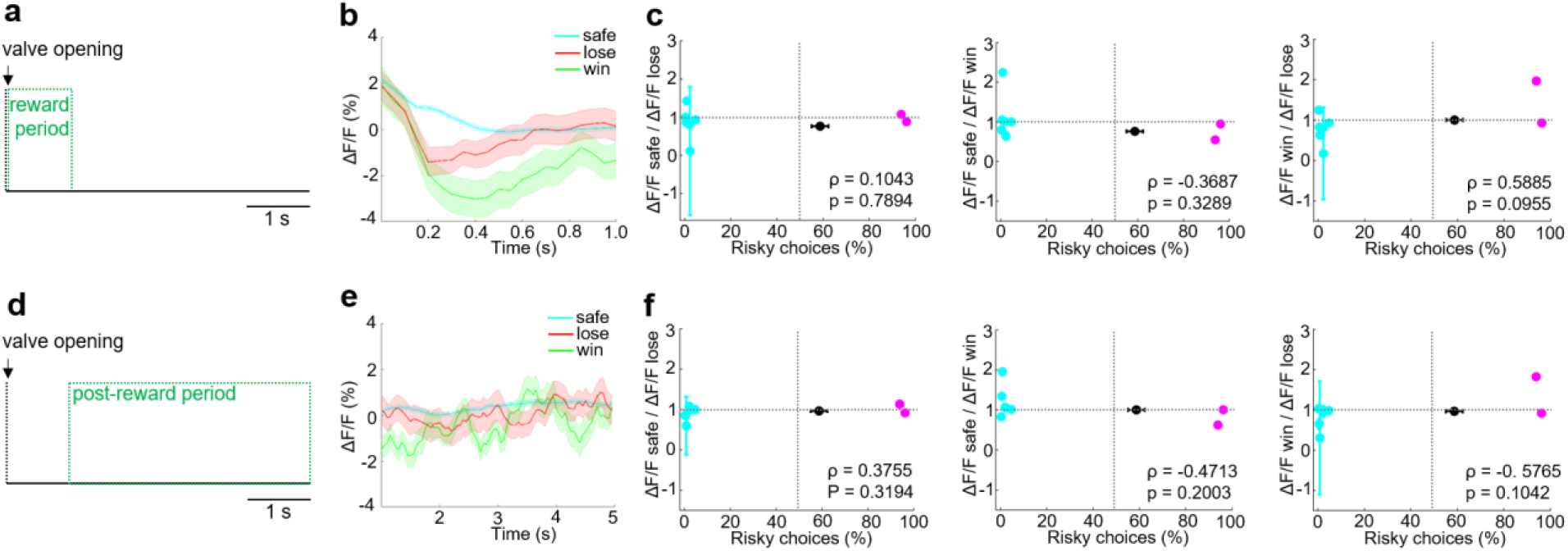
LHb fiber photometric signal in reward and post-reward period. **a**, Illustration of reward period used for analysis. **b**, LHb photometry example traces from one animal aligned to reward valve opening. **c**, Correlation of reward signal ratios and riskiness across animals. **d**, Illustration of post-reward period used for analysis. **e**, LHb photometry example traces for post-reward period (same animal as b). **f**, Correlation of post-reward signal ratios and riskiness across animals.

**Extended Data Fig 4.**
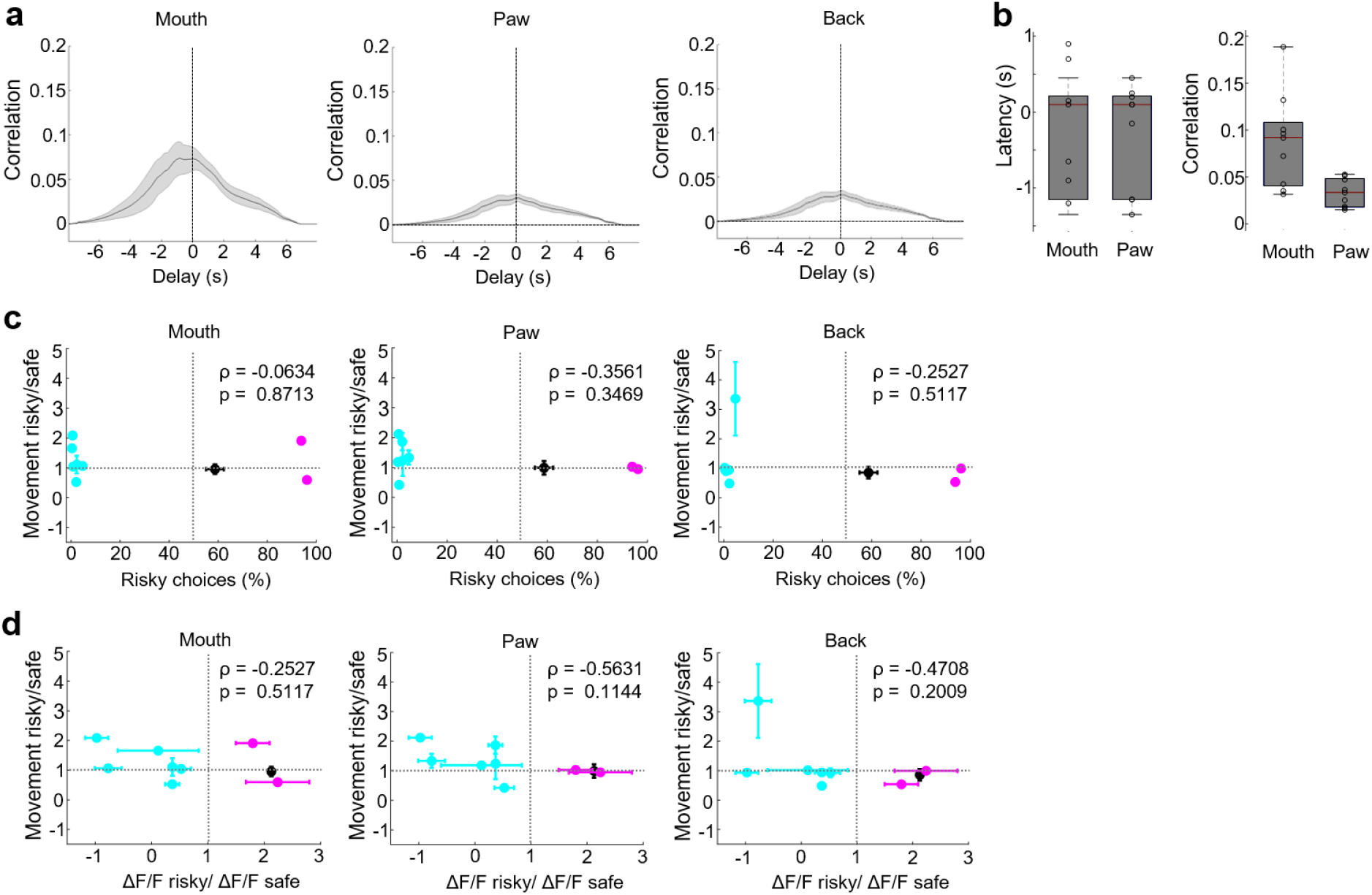
Divergence in LHb fiber photometric signal cannot be explained by body movement. **a**, Cross-correlation of body movement with bulk LHb signal. Data were obtained from the same animals as in Fig. 1F-K, and are presented as mean ± s.e.m. **b**, Quantification of cross correlation latency to peak and maximum amplitude across animals. **c**, Correlation of body movement and riskiness of the respective animal. **d**, Correlation of body movement and LHb photometry signal in the DP.

**Extended Data Fig 5.**
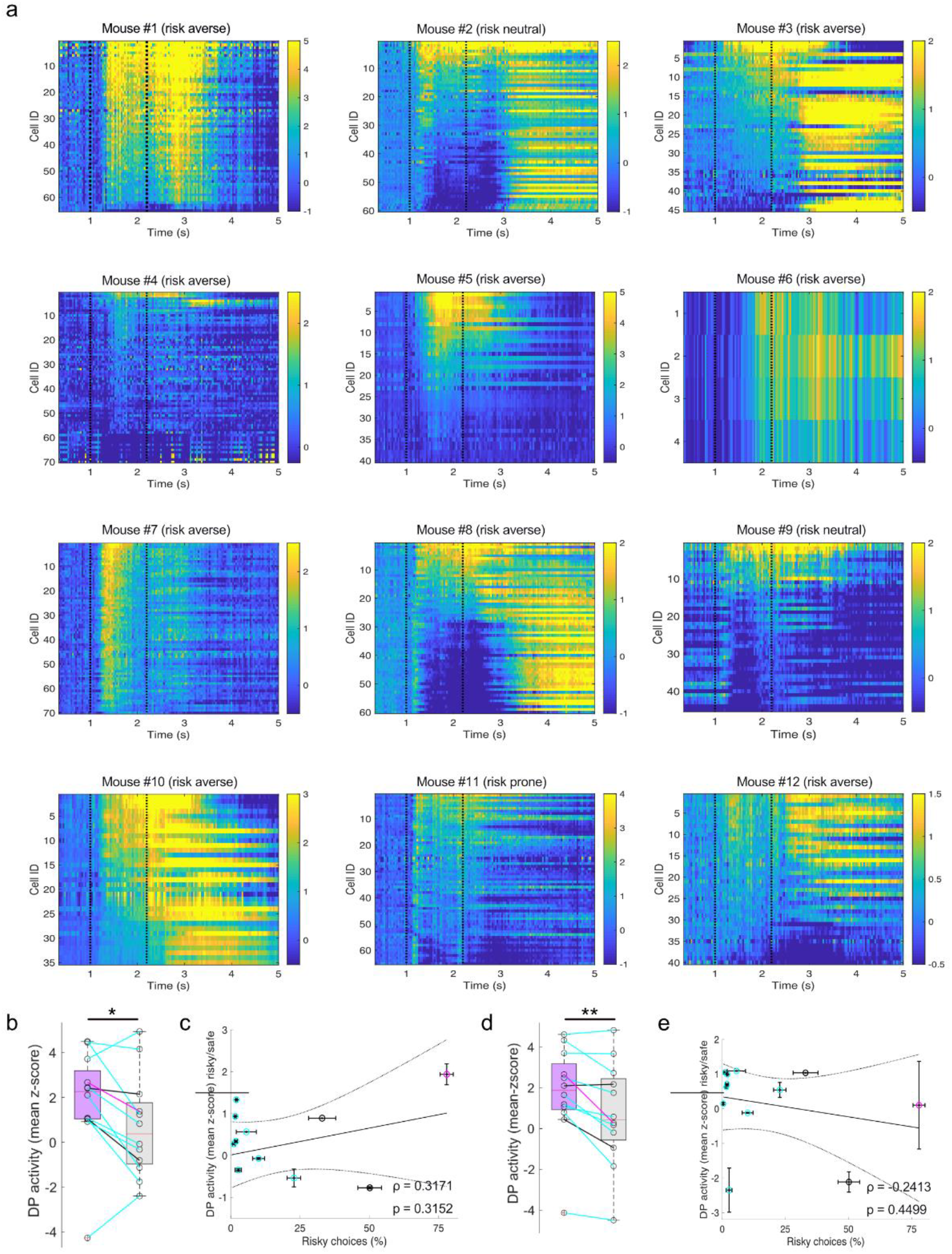
Inter-individual and bulk LHb two-photon data. **a**, Mean calcium transients (smoothed) for individual mice for all decisive trials sorted for mean activity in the DP. **b**, Quantification of two-photon bulk activity during the DP for present preferred (violet) and non-preferred (grey) choice trials. Line colors indicate individual risk preference (risk averse (cyan), risk neutral (black), risk prone (magenta), p=0.0425). Two-photon bulk data were achieved from drawing a single region of interest (ROI) containing all recorded cells across the entire field of view (FOV). **c**, Ratio of LHb ΔF/F integrals during the DP for present risky and safe choice bulk activity correlated with individual risk preference across sessions (n=12 mice). **d** and **e**, Same as b and c but trials were sorted for choice in the preceding trial (p=0.0049). Wilcoxon-signed rank test.

**Extended Data Fig. 6.**
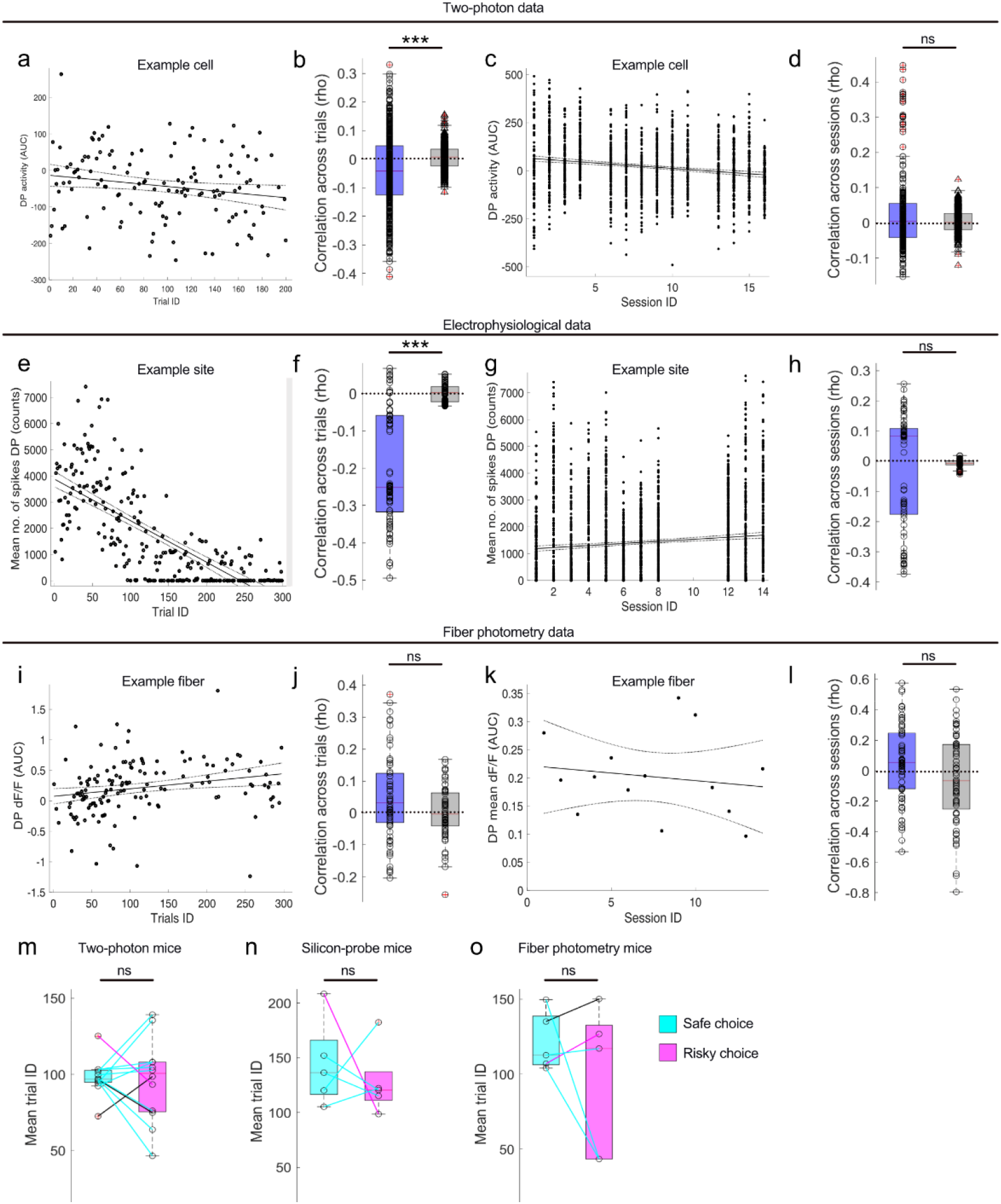
LHb DP activity attenuates over trials but not sessions and independent of trial choice. **a,** Single trial DP activity of LHb example cell (decisive trials only) across one example session. **b**, Quantification of mean single cell correlation with trial number (520 cells pooled across 12 mice; original (bue), shuffled (grey) data; p = 5.0193e-17). **c**, Activity of the same cell plotted across sessions. **d**, Quantification of across sessions (p = 0.1549). **e**-**h** Same as a-d but for multi-unit silicon probe recordings (90 recording sites across 5 mice (across trials (f): p = 3.1374e-14 ; across sessions (h): p = 0.27). i-l Same as a-d and e-h but for fiber photometric recordings of LHb bulk activity (n=5 mice; across trials (j): p = 0.0849; across sessions (l): p = 0.0543). Mean trial ID for safe (cyan) and risky (magenta) choices for two-photon (**m**; n=12 mice; p = 0.9697), silicon probe (**n**; n=5 mice; p = 0.6250), and fiber photometry (**o**; n=5 mice; p = 0.8125).Line colors indicate individual risk preference (risk averse (cyan), risk neutral (black), risk prone (magenta)). Wilcoxon signed rank test (paired). *** p<0.001

**Extended Data Fig 7.**
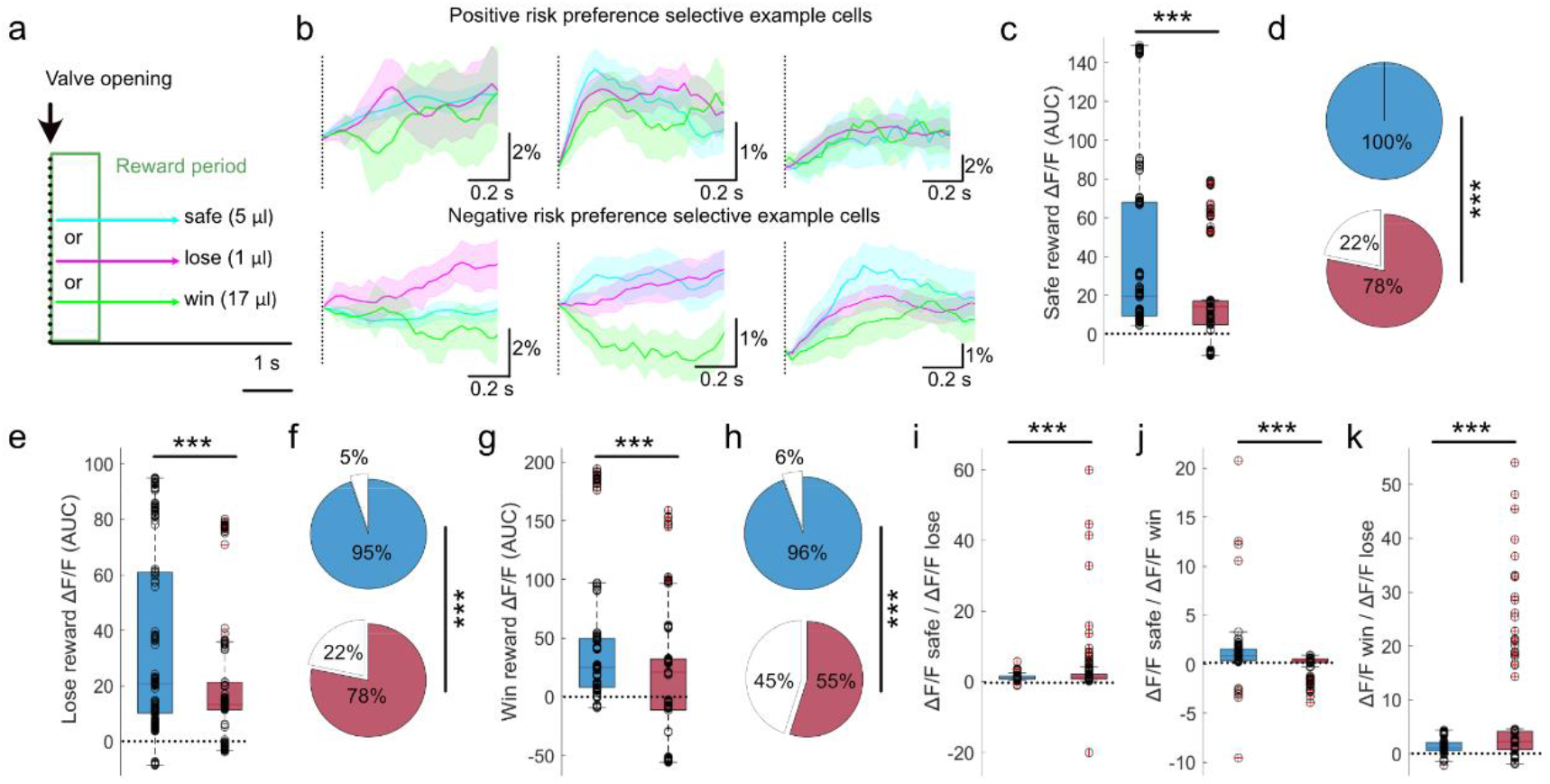
Positive and negative risk-preference selective cells (RPSCs) show divergent activity upon reward delivery. **a**, Illustration of reward period used for analysis. **b**, Example traces of three positive (top row) and three negative RPSCs (bottom row) aligned to reward delivery from three different animals (one positive and one negative cell for each mouse; from left to right: mouse #8, #2, and #13). Quantification of mean activity integrals (**c**; p= 4.29E-07)) and fraction of cells with positive values (**d**; p= 1.67E -06) for medium safe reward for positive (blue) and negative (red) RPSCs. Same as c and d but for small risky lose (**e**, p=5.26E -03 (activity);**f**, p= 1.18E -05 (fractions)) and for large risky win reward (**g**, p=2.08E -04 (activity); **h**, p= 1.02E -11 (fractions). Reward signal ratios for safe/lose (**i**; p=2.87E -04), safe/win (**j**; p=7.42E -19), and win/lose (**k**; p=9.86E -04).*** p<0.001, Wilcoxon rank sum test (activity), χ^2^-test (cell fractions), two-sampled Kolmogorov Smirnov test (activity ratios).

**Extended Data Fig 8.**
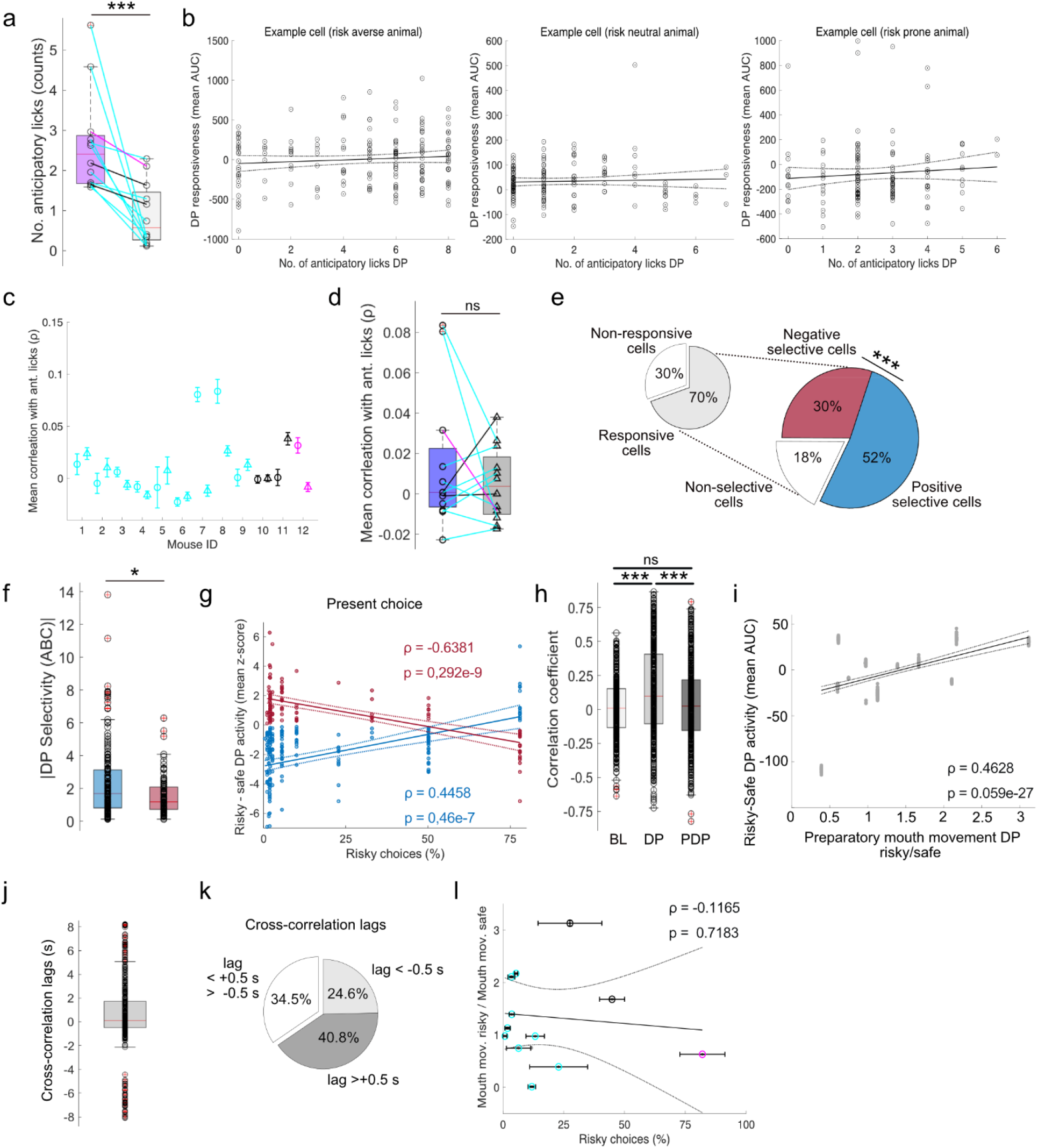
Anticipatory licking and preparatory mouth movement do not explain two-photon LHb single cell activity during the DP. **a**, Mean number of anticipatory licks during the DP for preferred (violet) and non-preferred choice trials. Line colors indicate individual risk preference (risk averse (cyan), risk neutral (black), risk prone (magenta); p=4.88E -04; Wilcoxon signed rank test). **b**, Correlation of mean DP activity with number of anticipatory licks for three example cells across three different mice. **c**, Mean correlation coefficient across all cells from individual mice for original (circle) and shuffled (triangle) data. **d**, Quantification of original and shuffled correlation coefficients (n=12 mice; p=0.8501; Wilcoxon signed rank test). Cell fractions (**e**, p=6.055E -06; χ^2^-test), RPSC selectivity (**f**, p=0.0182; Wilcoxon signed rank test) and correlation of RPSC DP activity difference between risky and safe choice with individual riskiness for present choices (**g**) calculated from trials without anticipatory licks. Positive RPSCs in blue negative RPSCs in red. **h**, Correlation of mean single cell activity with mouth movement during 1-s baseline period (BL), DP, and 1-s post-deliberation period (PDP; 483 cells from n=12 mice; Friedman-test; ***p<0.001; ns p>0,05). **i**, Correlation of preparatory mouth movement ratios (mean across trials) with mean activity of LHb neurons during DP (520 cells from n=12 mice). Cross-correlation lags of mean neuronal activity with mean preparatory mouth movement (**j**) and fraction of cells (**k**) based on 0.5 s-threshold (475 cells from n=12 mice). **l**, Correlation of individual riskiness with preparatory mouth movement ratios (n=12 mice).

**Extended Data Fig. 9.**
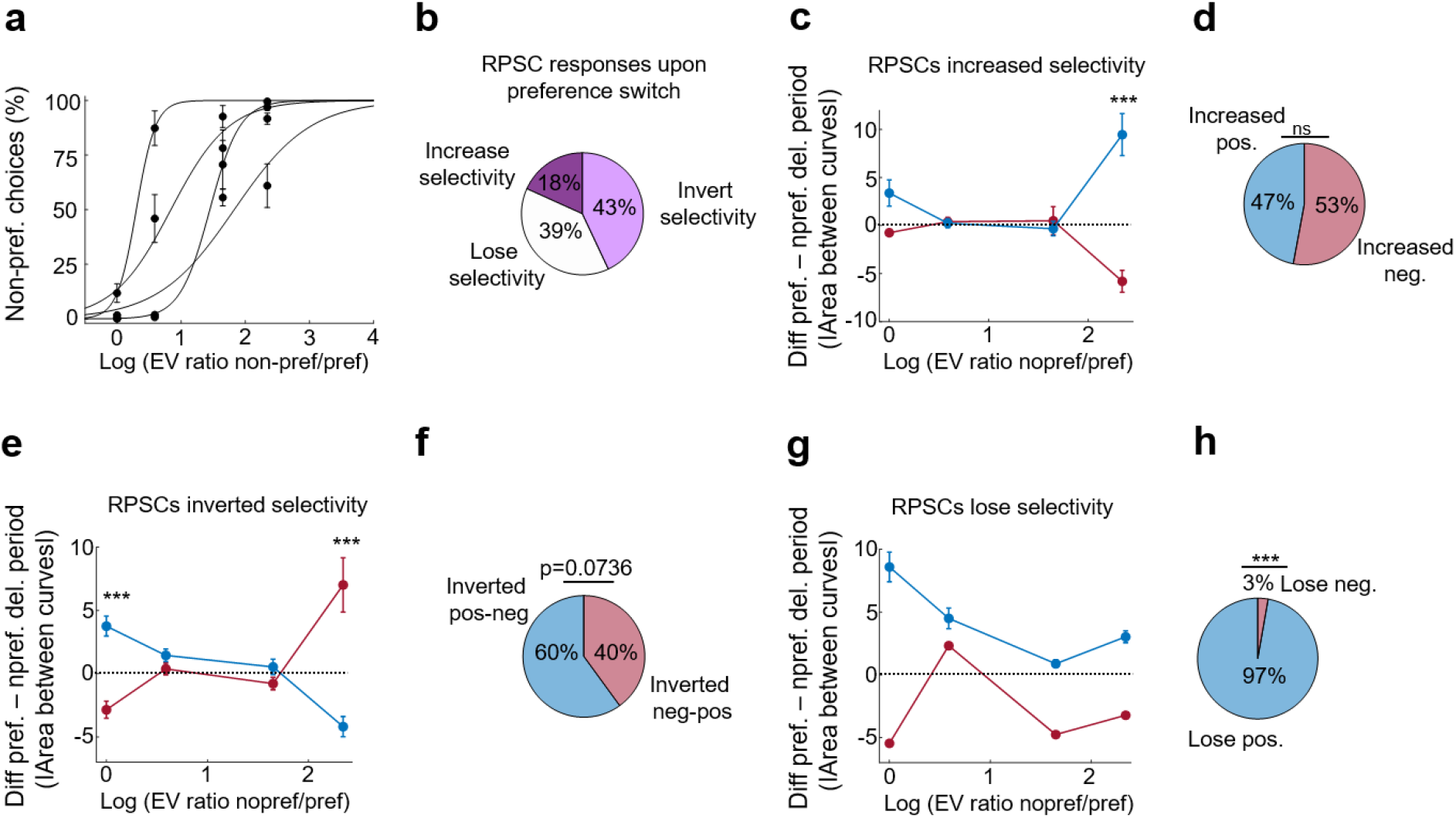
LHb RPSCs adapt their selectivity DP with changing preference. **a**, Fraction of non-preferred option choices with changing expected value ratio (n = 4 mice). Every dark circle represents 3-5 sessions with 200 free choice trials each. Sigmoid fit. **b**, Fractions of LHb RPSCs tracked across sessions depending on their adaptation to preference change (baseline sessions compared to last highest ratio sessions). Positive (blue) and negative (red) LHb RPSCs (selectivity and fraction of cells) depending on their responses: Cells either increased (**c**,**d**), inverted (**e**, **f**), or lost (**g**, **h**) their selectivity.

**Extended Data Fig. 10.**
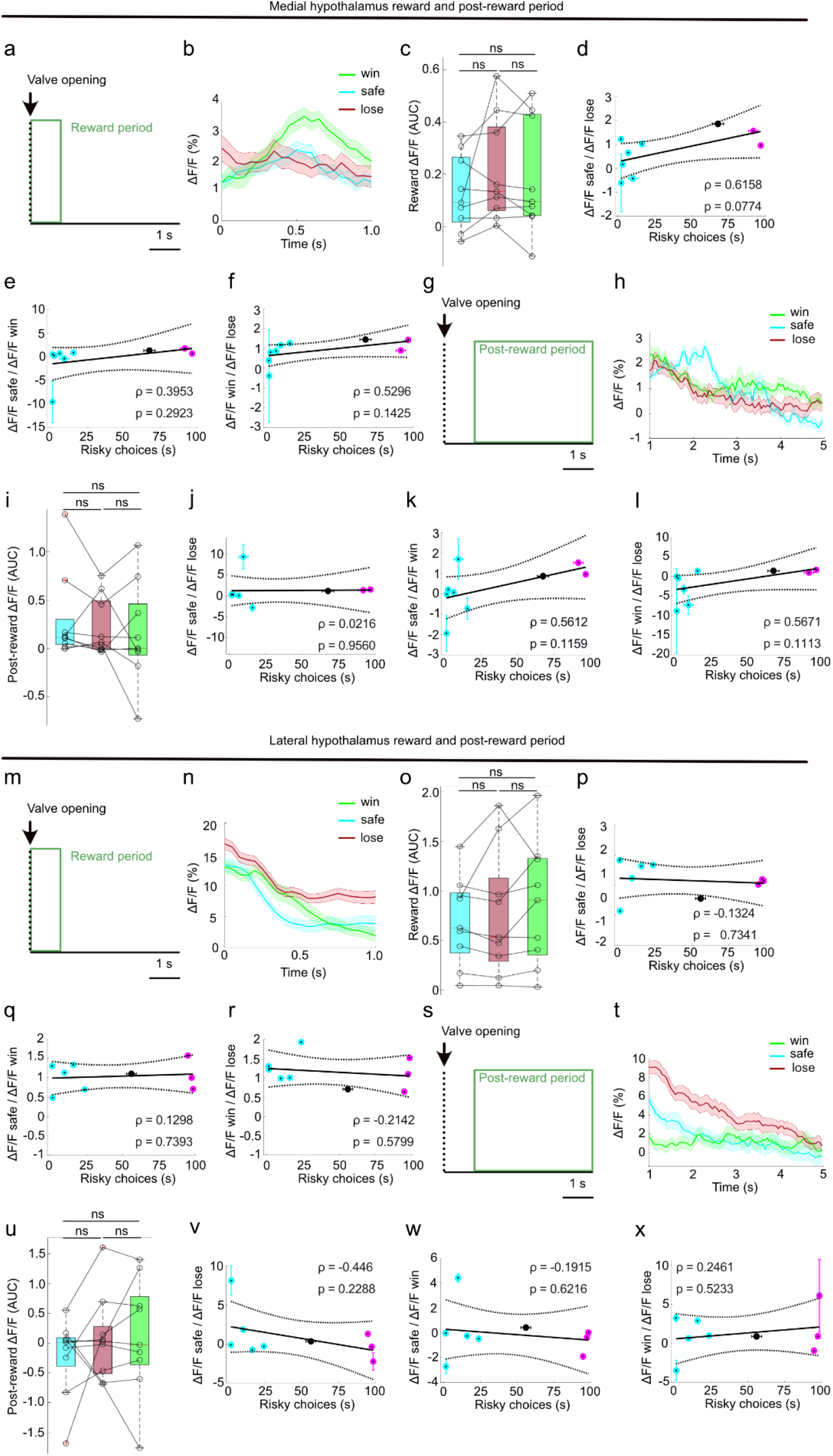
LH and MH fiber photometric signal in reward and post-reward period. **a**, Illustration of reward period used for analysis. **b**, MH photometry example traces from one animal aligned to reward delivery. **c**, Quantification of MH population activity integrals aligned to reward delivery for medium safe (cyan), small risky lose (red), and large risky win (green) reward outcomes (p=0.2199; n=9 mice). Correlation of reward signal ratios and riskiness for safe/lose (**d**), safe/win (**e**), and win/lose (**f**). Same as a-f but for post-reward period (1-5 s post reward period; **g**-**l**; p=0.4305). Same as a-l but for LH population activity (n=9 mice; **m**-**x**; reward period p = 0.3228; post-reward period p=0.4959). ns: p>0.05; repeated measure ANOVA with Bonferroni’s multiple comparison test.

**Supplementary Fig. 1.**
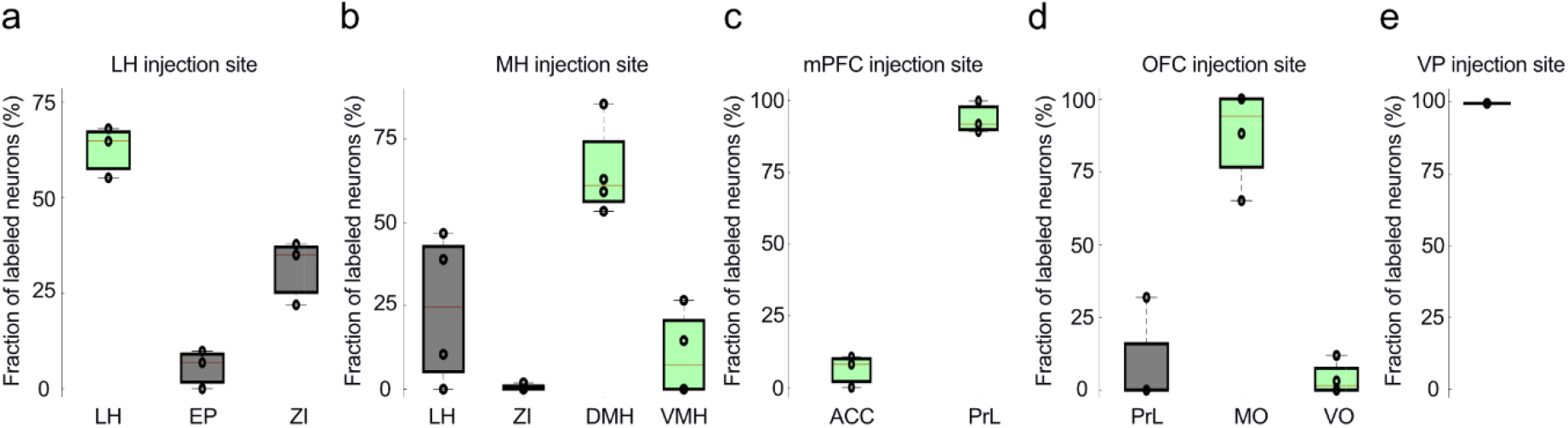
On- and off-target labelling of anterograde tracing. **a**, Fraction of labeled neurons from viral injections targeting the lateral hypothalamus (LH; entopeduncular nucleus (EP); zona incerta (ZI); **a**), medial hypothalamus (MH;dorsomedial hypothalamus (DMH), ventromedial hypothalamus (VMH); **b**), medial prefrontal cortex (mPFC; anterior cingulate cortex (ACC); prelimbic cortex (PrL); **c**), orbital frontal coretex (OFC; medial orbital frontal cortex (MO); ventral orbital frontal cortex (VO); **d**), and ventral pallidum (VP).

**Supplementary Fig. 2.**
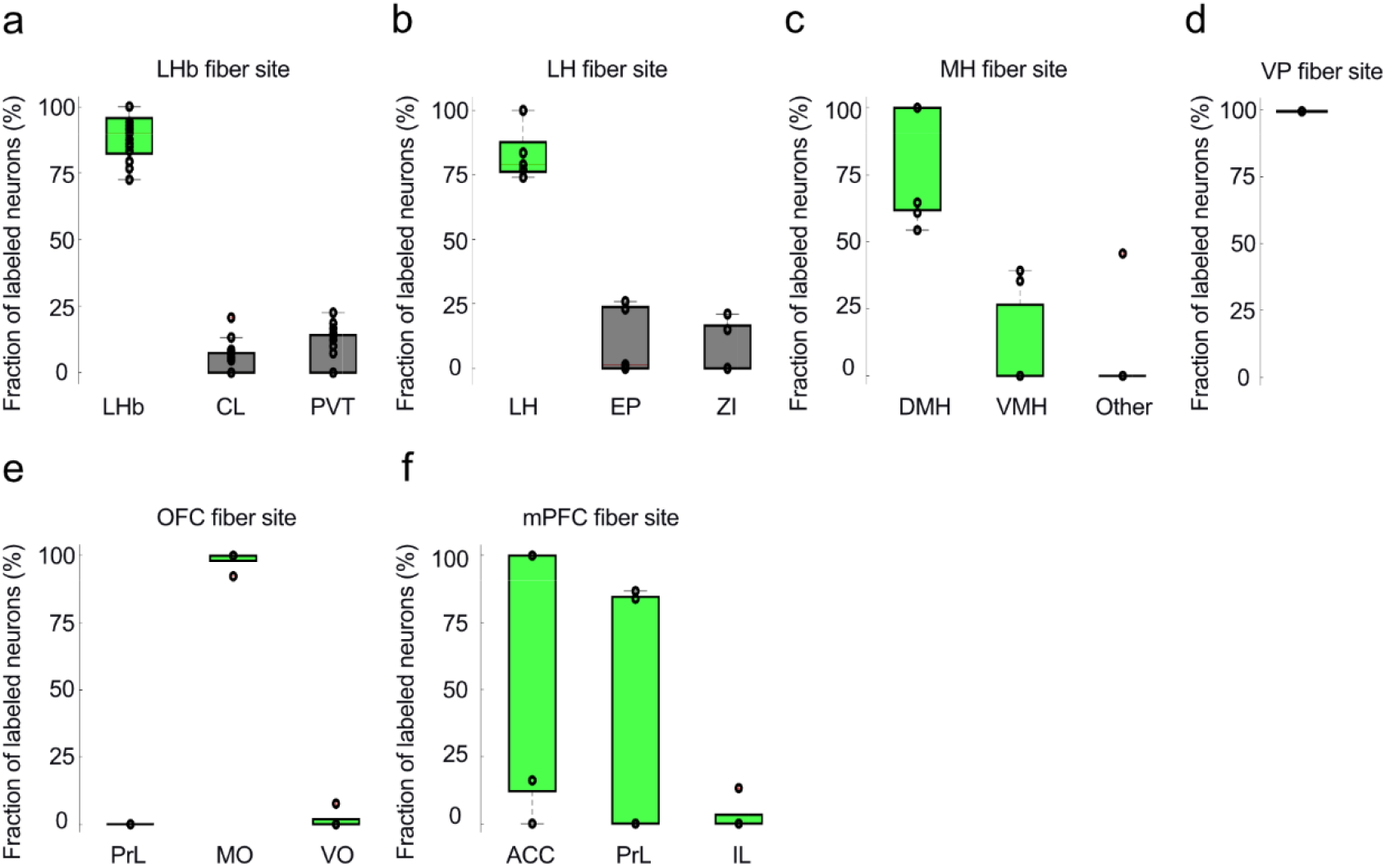
On- and off-target labelling of multi-fiber recordings. Quantification of GCaMP6f-labeled somata within the maximum range of fiber fluorescence detection for LHb (**a**; 400 μm; n=17), LH (**b**; 400 μm), MH (**c**; 200 μm), VP (**d**; 400 μm), OFC (**e**; 200 μm), and mPFC (**f**; 400 μm).

**Supplementary Fig. 3.**
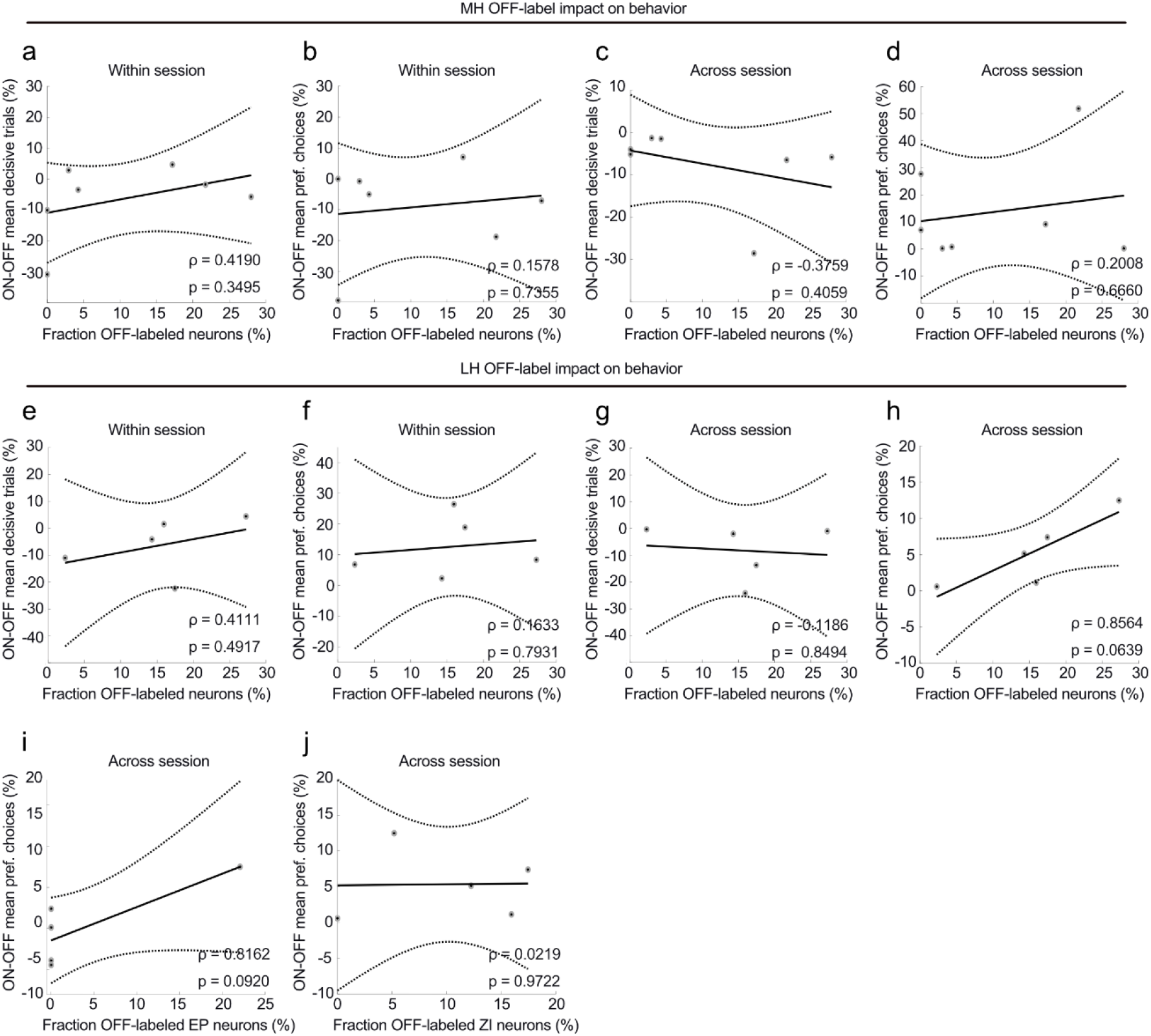
Off-target effect on decisiveness and choice preference of retrograde optogenetic perturbation. Impact of all off-target labeled (ArchT-expression) MH adjacent neurons on within (**a**, **b**) and across session parameters (**c**, **d**; n=7 mice). Same as a-d but for LH adjacent neurons (**e**-**h**; n=5 mice). Impact of LH adjacent, off-target labeled EP (**i**) and ZI (**j**) neurons on across session preference level.

**Supplementary Fig. 4.**
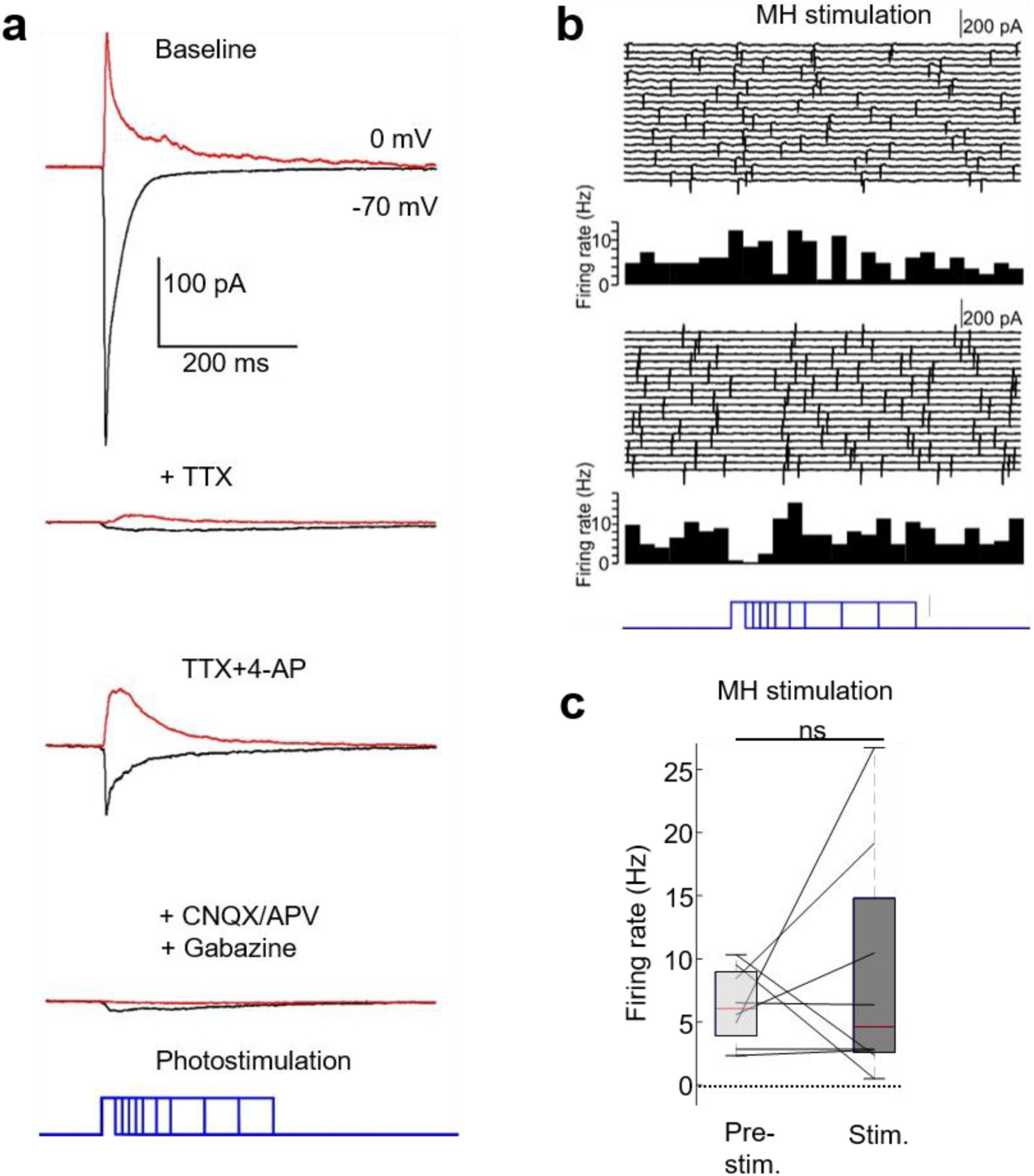
Optogenetic stimulation of hypothalamic terminals in acute LHb containing brain slices. **a**, Example traces of MH connected LHb cell show EPSCs (black) and IPSC (red) which are sensitive to sodium channel blocker tetrodotoxin (TTX). EPSCs and IPSCs are both monosynaptic connections, as they are maintained in TTX/4-AP, and sensitive to glutamate receptor antagonists (CNQX/APV), and GABA_A_-receptor antagonist Gabazine, respectively. Example spiking traces of two MH- connected LHb neurons (**b**; band-pass filtered at 1 and 1000 Hz) and quantification (**c**) of all LHb neurons tested for MH connectivity (loose seal recordings, 7 cells from 4 mice; Wilcoxon signed rank test; p = 0.5625) from two 20 ms bins before (pre-stim.) and during (stim.) optogenetic stimulation of MH axons.

**Extended Data Table 1.**
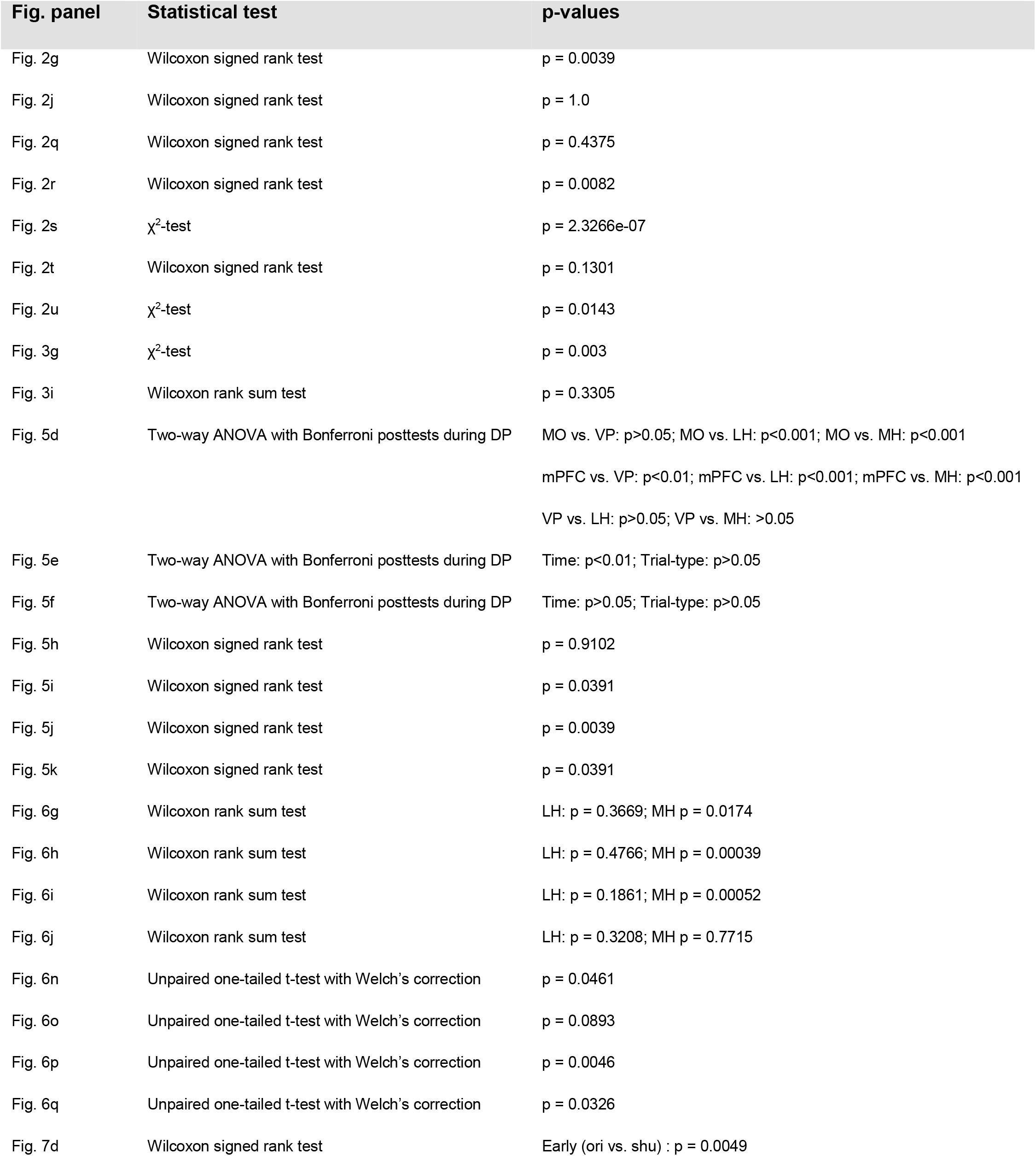
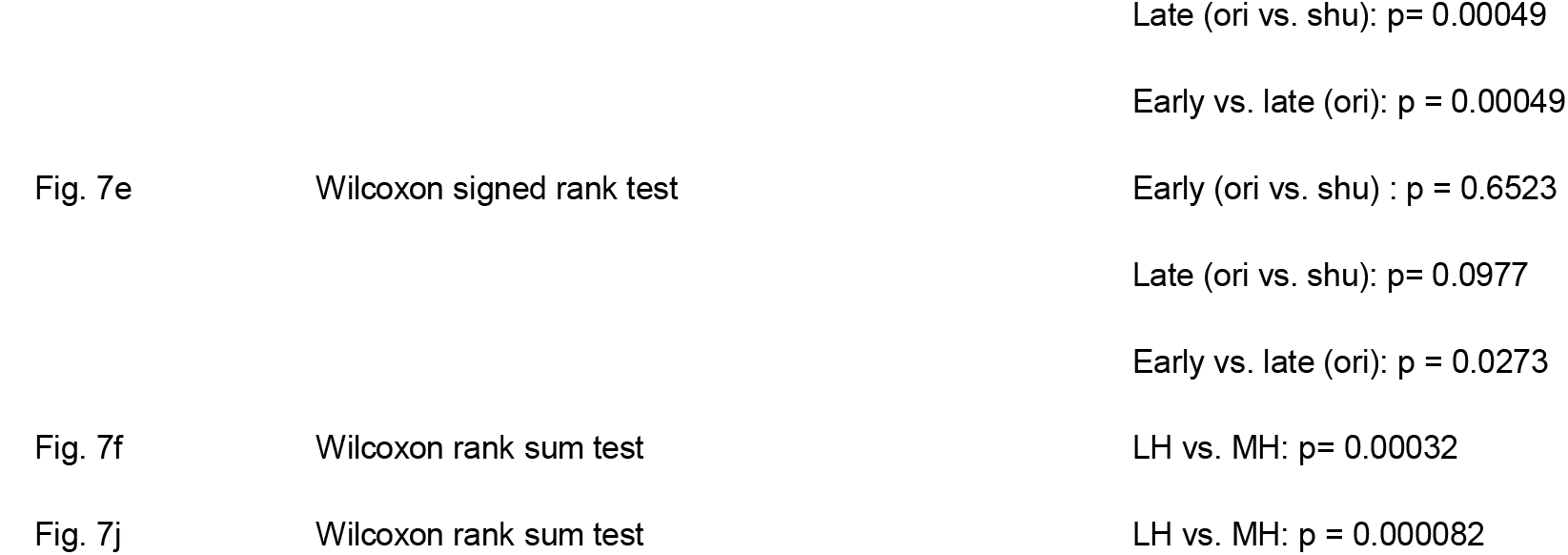
Statistical tests and exact p-values.

**Extended Data Table 2.**
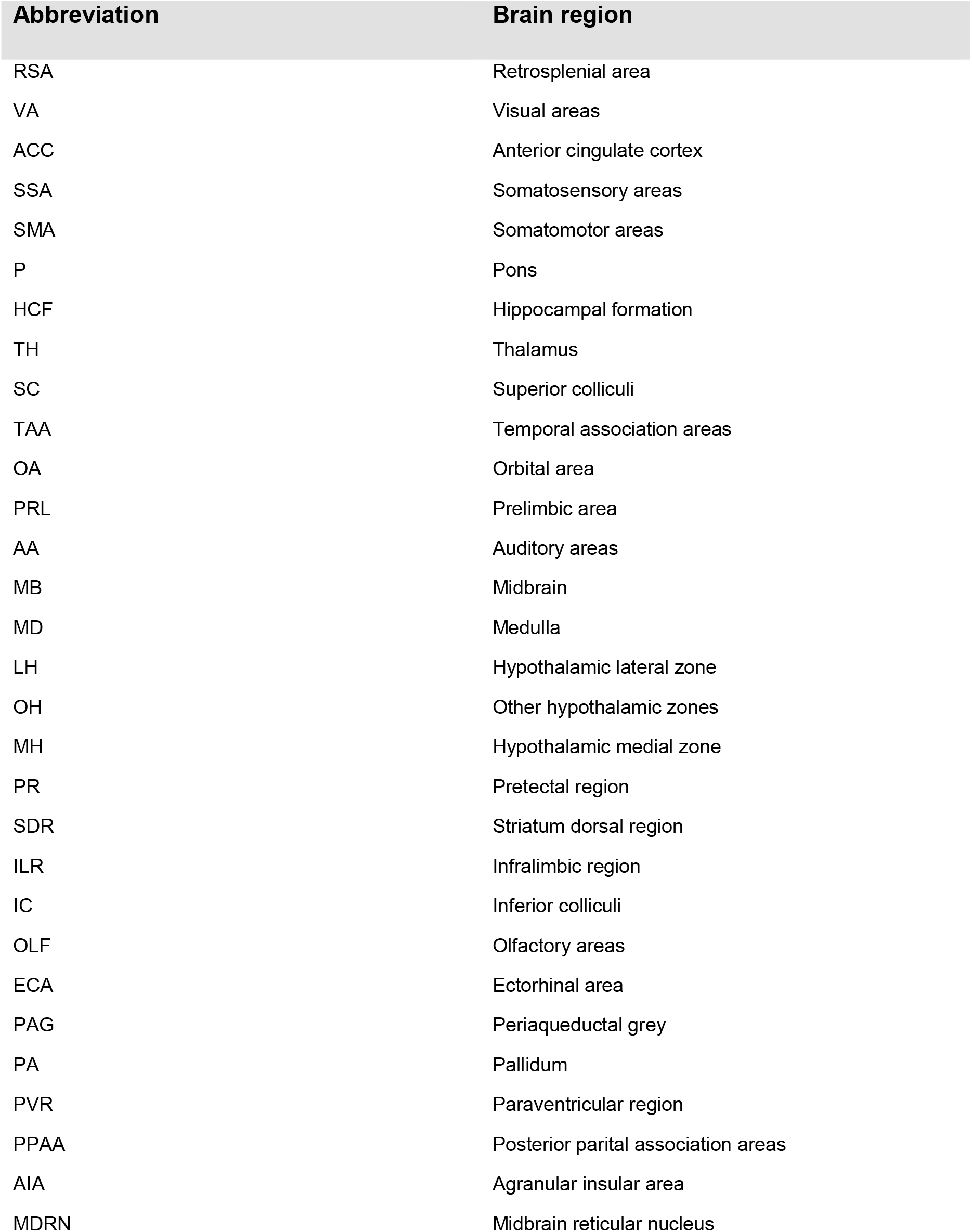
Abbreviation of retrogradely labelled LHb input regions.

| **Abbreviation** | **Brain region** |
| --- | --- |
| RSA | Retrosplenial area |
| VA | Visual areas |
| ACC | Anterior cingulate cortex |
| SSA | Somatosensory areas |
| SMA | Somatomotor areas |
| P | Pons |
| HCF | Hippocampal formation |
| TH | Thalamus |
| SC | Superior colliculi |
| TAA | Temporal association areas |
| OA | Orbital area |
| PRL | Prelimbic area |
| AA | Auditory areas |
| MB | Midbrain |
| MD | Medulla |
| LH | Hypothalamic lateral zone |
| OH | Other hypothalamic zones |
| MH | Hypothalamic medial zone |
| PR | Pretectal region |
| SDR | Striatum dorsal region |
| ILR | Infralimbic region |
| IC | Inferior colliculi |
| OLF | Olfactory areas |
| ECA | Ectorhinal area |
| PAG | Periaqueductal grey |
| PA | Pallidum |
| PVR | Paraventricular region |
| PPAA | Posterior parital association areas |
| AIA | Agranular insular area |
| MDRN | Midbrain reticular nucleus |

